# Resolving Gene-Metabolism Links through Multi-Metabolite mQTL Commonality Analysis

**DOI:** 10.64898/2026.11.03.763898

**Authors:** Leah Rosental, Yair Yehoshua Zach, Taly Lapidot-Cohen, Simy Weil, Claire Tabachnik, Ezra Sternlicht, Tali Niger, Anna Borovikova-Sheinker, Sayantan Panda, Ilana Rogachev, Nir Shahaf, Bo Ouyang, Asaph Aharoni, Shuchao Dong, Nir Sade, Yariv Brotman

**Affiliations:** Department of Life Sciences, Ben Gurion University of the Negev, Beersheva, Israel; School of Plant Sciences and Food Security, Institute for Cereal Crops Research, Tel Aviv University, Tel Aviv, 69978 Israel; Department of Plant and Environmental Sciences, Weizmann Institute of Science, Rehovot, Israel; National Key Laboratory for Germplasm Innovation and Utilization of Horticultural Crops, Huazhong Agricultural University, Wuhan 430070, China; Institute of Vegetable Crop, Jiangsu Key Laboratory for Horticultural Crops Genetic Improvement, Jiangsu Academy of Agricultural Sciences, Nanjing 210014, China

## Abstract

High-throughput metabolomics detects so many compounds that most studies use only a fraction of them, focusing on the few metabolites with an annotated structure or substructure. Metabolite genome-wide association studies of specialized metabolism detect many metabolite quantitative trait loci (mQTLs), but most associated metabolite features remain unannotated, limiting gene discovery. In leaves of 362 tomato accessions under control and water-deficit conditions, only 13 of the 826 metabolite features associated with 77 reproducible mQTLs were annotated with good confidence. To study the mQTLs with the largest impact, we developed commonality analysis, an approach that does not depend on well-studied pathways.

Commonality analysis uses biologically defined groups, such as the features associated with one mQTL, to infer relatedness between metabolites. By identifying the fragments, neutral losses or mass transformations that are common in a group, it links the group to a substructure, pathway or reaction, and can extend annotation within the group. Most high-impact mQTLs had a significantly distinct fragment commonality, whose outstanding fragments suggested the pathway involved and its candidate gene. We demonstrate the approach in three representative mQTLs: (i) common fragments pointed to a sinapoyl-hexose pathway and a serine carboxypeptidase-like candidate acyltransferase; (ii) common fragments marked the features associated with a purine permease as nitrogen-containing, which isotope labeling then supported; (iii) a common 2.02 Da transformation pointed to a double-bond reduction and a putative reductase. Overexpression of each gene reproduced the commonality patterns, supporting the candidate genes and showing that the approach also applies to experimentally defined metabolite groups.

## Introduction

Plants produce an extraordinary diversity of metabolites that support growth, development, defense, and environmental adaptation. Tomato (*Solanum lycopersicum*) provides an excellent system for studying genetic variation and crop metabolism. Extensive genomic resources and germplasm collections^1–4^ have facilitated the genetic dissection of traits related to domestication, fruit quality, flavor, and metabolite composition^5–9^. However, most tomato metabolomic studies have focused on fruit-associated metabolites, whereas the genetic regulation of the leaf metabolome remains largely unexplored^10^, despite the central role of leaves in photosynthesis, productivity, and environmental response^11,12^.

Using the natural variation of metabolite abundances as phenotypic traits in metabolite genome-wide association studies (mGWAS) enables the identification of metabolite quantitative trait loci (mQTLs) in the genome and facilitates the discovery of metabolic genes, pathways, and regulatory networks^13–15^. Environmental perturbation expands the repertoire of metabolites produced and consequently the power of mGWAS to reveal mQTLs that are weak or absent under other conditions, as well as those stable in both^16–20^. However, moving from statistical association to biochemical interpretation usually requires information on the identity of the metabolites involved.

Despite advances in metabolite detection by liquid chromatography-mass spectrometry (LC-MS), most specialized metabolites remain unannotated^21,22^. Current annotation approaches commonly match MS/MS (or MS^2^) spectra to open-source reference libraries such as MassBank and GNPS^23,24^ or vendor-specific libraries such as mzCloud and METLIN. While effective, direct library matching excludes the identification of metabolites missing from reference libraries. To expand the annotation of metabolites, multiple approaches integrate the metabolite information obtained by LC-MS/MS in various ways. For example, Feature-Based Molecular Networking (FBMN) links metabolites according to MS^2^ spectral similarity, enabling propagation of substructure annotations within related chemical families^24,25^. The SIRIUS software uses MS^2^ spectra to predict chemical fingerprints of molecular substructures and molecule structures^26^. The MS2LDA algorithm identifies recurring fragment and neutral-loss motifs that facilitate substructure annotation of unknown compounds^27^. Although the number of approaches and *in silico* tools for expanding metabolite annotation is increasing, their accuracy and suitability for annotating *de novo* metabolites and chemical families are still limited. For example, a systematic evaluation of fragment-ion annotations in three major *in silico* MS^2^ libraries (HMDB, METLIN, and mzCloud) revealed substantial inaccuracies, highlighting the enduring limitations of current annotation approaches^28^.

Beyond spectral similarity, biochemical relationships between metabolites can provide an additional source of information. Candidate substrate-product pair (CSPP) networks identify metabolites separated by characteristic mass differences corresponding to enzymatic transformations^29^, which can be used in combination with molecular networking to identify recurrent biochemical modifications across metabolomes^30^. Tools such as MetNet^31^, NetID^32^ and MetDNA^33^ use biologically relevant mass differences and metabolic connectivity to improve metabolite interpretation.

In the context of mGWAS, several methods have begun to integrate biochemical information directly into genetic analyses. For example, chemical-tag-based semi-annotated metabolomics demonstrated that recurring modifications can support candidate-gene discovery and pathway reconstruction without requiring full metabolite identification^34^. Shortest reactional distance (SRD) analysis, which is based on KEGG reactions, incorporated metabolic pathway information to prioritize gene-metabolite relationships beyond conventional significance thresholds^35^. Together, these studies illustrate the growing effort to move from statistical associations toward mechanistic interpretation of mGWAS datasets. Nevertheless, most studies still rely on annotated structures to guide the selection of genes for functional characterization^36–38^. Considering that only ∼1% of plant specialized metabolites have been annotated, a recent perspective argued that the next major advance in plant metabolomics will require overcoming the strong bias toward the limited number of annotated metabolite families^39^. Further, these methods do not systematically exploit the co-association of multiple metabolite features as an information source in their analysis.

Here, we present a large-scale analysis of leaf metabolism in a genetically diverse tomato population grown under control and prolonged water-deficit conditions. We provide a resource comprising transcriptomes, hundreds of chromatograms, data-dependent tandem mass spectra (ddMS^2^), curated metabolite annotations and mQTL associations.

To confront the scale and complexity of the data and to study metabolic gene regulation in an unbiased manner, we developed commonality analysis, which extracts shared mass descriptors (fragment ions, neutral losses and transformations) from the features associated with each mQTL. We show that most high-impact mQTLs, those with enough features to statistically test the mass descriptors, have a significantly distinct commonality, which helps identify candidate pathways, reactions, and genes underlying metabolic variation. We present three examples of how this strategy improves metabolite annotation and enables identification and functional characterization of genes underlying mQTLs.

## Results

To investigate the genetic regulation of tomato leaf metabolism, a diverse tomato collection was grown in a greenhouse experiment. The “TS” collection includes 362 accessions with representatives from *S. pimpinellifolium, S. lycopersicum* var. cerasiforme and diverse *S. lycopersicum* tomato varieties including many heirloom, transitional and modern cultivars^2,6^. Because water stress substantially alters leaf metabolism^40–43^, we evaluated the population under two conditions: typical agricultural production conditions (control) and reduced water availability (water deficit; WD). Comprehensive evaluations of gene expression and metabolism were integrated with high-density genomic polymorphism data^2^ to enable thorough investigation (Fig. 1A).

**Figure 1:**
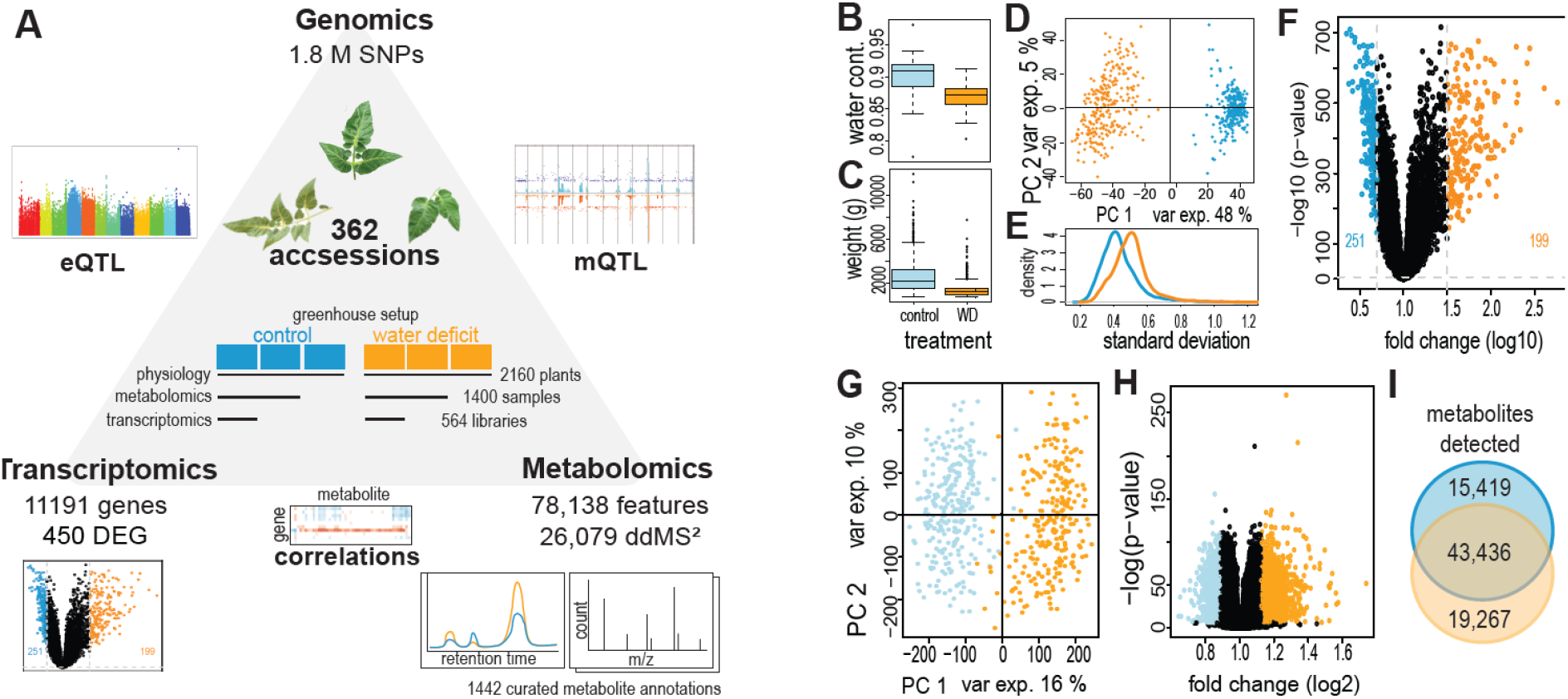
Experiment overview and stress-induced variation. A. Overview of experiment setup, datasets produced and the main analysis and results to which they led. B. Leaf water content, as the ratio of leaf weight lost after full oven drying to the fresh leaf weight, in leaves collected three weeks after treatment onset from control (light blue) and water deficit (WD; orange) treated plants. N_control_=155, N_WD_=54. C. The fresh vegetative weight of whole plants measured after last fruit harvest. N_control_=656, N_WD_=589. D. Global effect of WD on leaf gene expression. PCA based on relative gene expression of leaves collected from control (blue) and water deficit (dark orange) treated plants. E. Distribution of variation (standard deviation) in the expression of each gene across the population in either growth condition. F. Gene expression in response to WD. Volcano plot displaying the significance of a t- test comparing transcript levels between water deficit and control conditions to the fold change (after log10 transformation) of the same comparison. Transcripts with significant increase (p-value <α=0.05, Bonferroni correction) and fold change exceeding double expression in water deficit (dark orange) or control (blue) are colored. G. Global effect of WD on metabolic profile. PCA based on metabolic profiles of leaves collected from control (light blue) and water deficit (orange) treated plants with metabolite features detected in both positive and negative ionization modes. H. Feature changes in response to WD. Volcano plot displaying the significance of a paired t-test comparing feature levels of each accession between WD and control conditions to the median fold change (after log2 transformation). Features with significant increase (p-value <α=0.05, Bonferroni correction) and fold change exceeding two-fold expression in water deficit (orange) or control (light blue) are colored. I. The number of features detected exclusively in control (blue circle), water deficit (orange circle) or in both (overlap).

### Gene expression and metabolic variation under two conditions

We first set out to evaluate the severity of the imposed stress and the changes it caused. The applied WD caused a significant and substantial drop in leaf water content (Fig. 1B), but allowed plant survival, albeit with strongly reduced vegetative productivity (Fig. 1C) by the end of the growing season. Transcriptomic analysis quantified relative expression of 11,191 genes in 539 samples (Supp. Table 1). As expected, a global shift in the gene expression pattern between control and WD plants was found (Fig. 1D), explaining 48% of the variation in the first principal component. Most genes differed significantly but mildly (low population-wide fold change) between conditions. Still, some genes changed significantly and substantially: 251 (2.2%) genes decreased (fold change < 0.5) and 199 (1.7%) increased (fold change > 2) under WD (Fig. 1F). Population-wide variance in gene expression was greater among WD samples, as can be seen in the wider spread of samples along the first and second principal components and in the increase in expression variation of individual genes within the population (Fig. 1E). Gene Ontology (GO) analysis revealed that genes increased in WD were enriched in secondary metabolite biosynthetic process (GO:0044550) genes, whereas general metabolic process (GO:0008152) and other central metabolic process related GO terms were depleted (Supp. Figure 1; Supp. Table 2).

Leaf metabolomic profiling was conducted using an untargeted LC-MS method which detects polar and semi-polar metabolites. Over 100,000 metabolite features (peaks with a unique mass-to-charge ratio (m/z) and retention time (RT) after isotope clustering, hereafter features) were detected in positive and negative ionization modes combined. After applying quality control filters, 78,138 were retained for the following analyses.

Growth under WD greatly altered the metabolic composition of the leaves (Fig. 1G). Accordingly, the vast majority of features were significantly increased or decreased in WD. Although the significant changes were often mild, over 3.5% of features changed by more than 2-fold between the conditions (Fig. 1H). In addition, we detected 15,419 (19%) features exclusively in control conditions and 19,267 (24%) in WD, highlighting the benefit of including multiple conditions for a comprehensive analysis (Fig. 1I).

### Leaf metabolite annotations

One major bottleneck in metabolomics is assigning structural information to detected features^44,45^. We therefore combined multiple annotation methods to maximize annotation coverage (Supp. Table 3).

Thirty diverse samples from our GWAS experiment were chosen for comprehensive feature ddMS^2^ fragmentation, leading to 13,781 ddMS^2^ spectra obtained in positive ionization mode and 12,298 in negative mode. We constructed a feature-based molecular network (FBMN) from the ddMS^2^ spectra and matched them to the GNPS^46^ library (Supp. Figure 2). The network included 548 well-matched library annotations, but many matched multiple features at different RTs. Next, we searched several public and in-house libraries using Compound Discoverer (Thermo Fisher Scientific). High-confidence matches to the in-house run library (based on MSMLS, Mass Spectrometry Metabolite Library of Standards, Sigma-Aldrich) identified 81 features as 64 unique metabolites. External library searches matched 458 features to metabolite MS^2^ spectra. Additional comparisons to published LC-MS based annotations gave putative annotations to 161 features, but at lower confidence because they were based on fragments co-locating in MS^1^ rather than on ddMS^2^ matches. The same set of 30 samples was analyzed with WeizMass^47^ using its optimized LC-MS method^9^, yielding high-confidence metabolite annotations for 122 features. Each method contributed some new annotations, but as expected many features had redundant identifications from multiple sources. Overall, we curated metabolite annotations for 1,442 features (Supp. Table 3), with varying annotation confidence levels^48^, which were later used to interpret the mGWAS results.

### Overview of mQTLs detected in control and in water deficit conditions

The abundance of each feature was used as a phenotype for GWAS^2^, with two independent repetitions for each condition (control and WD). Adjacent genomic associations of the same feature within a specific window of 100 kb were collapsed and defined as a single mQTL. Features associated with overlapping genomic regions were assigned to the same mQTL and together form its mQTL feature group.

In total, 719 unique mQTLs were found (Fig. 2A; Supp. Table 4). In control conditions 410 mQTLs were found, with 192 repeatedly confirmed in the second control replicate. For WD 401 mQTLs were found, and 186 were confirmed by repeated association in WD. Many mQTLs were detected only once. These tended to have very few associated features, reflecting the risk of random sporadic associations in such high-throughput analysis. We therefore considered only mQTLs for which the feature association was reproduced in the second repetition. We found that 36 mQTLs were constitutive with consistent feature associations in both repetitions of the two conditions, while 13 or 16 mQTLs had features repeatedly associated only in WD or in control, respectively. Interestingly, 12 mQTLs had features repeatedly associated in WD and other features repeatedly associated in control. This could be caused by mistaken co-association because of closely linked polymorphisms or by activity on features present exclusively in either condition. Together, we detected 77 reproducible mQTLs. Again, the condition-specific mQTLs show the benefit of investigating different conditions for obtaining a more complete picture of genetic control of leaf metabolism.

**Figure 2:**
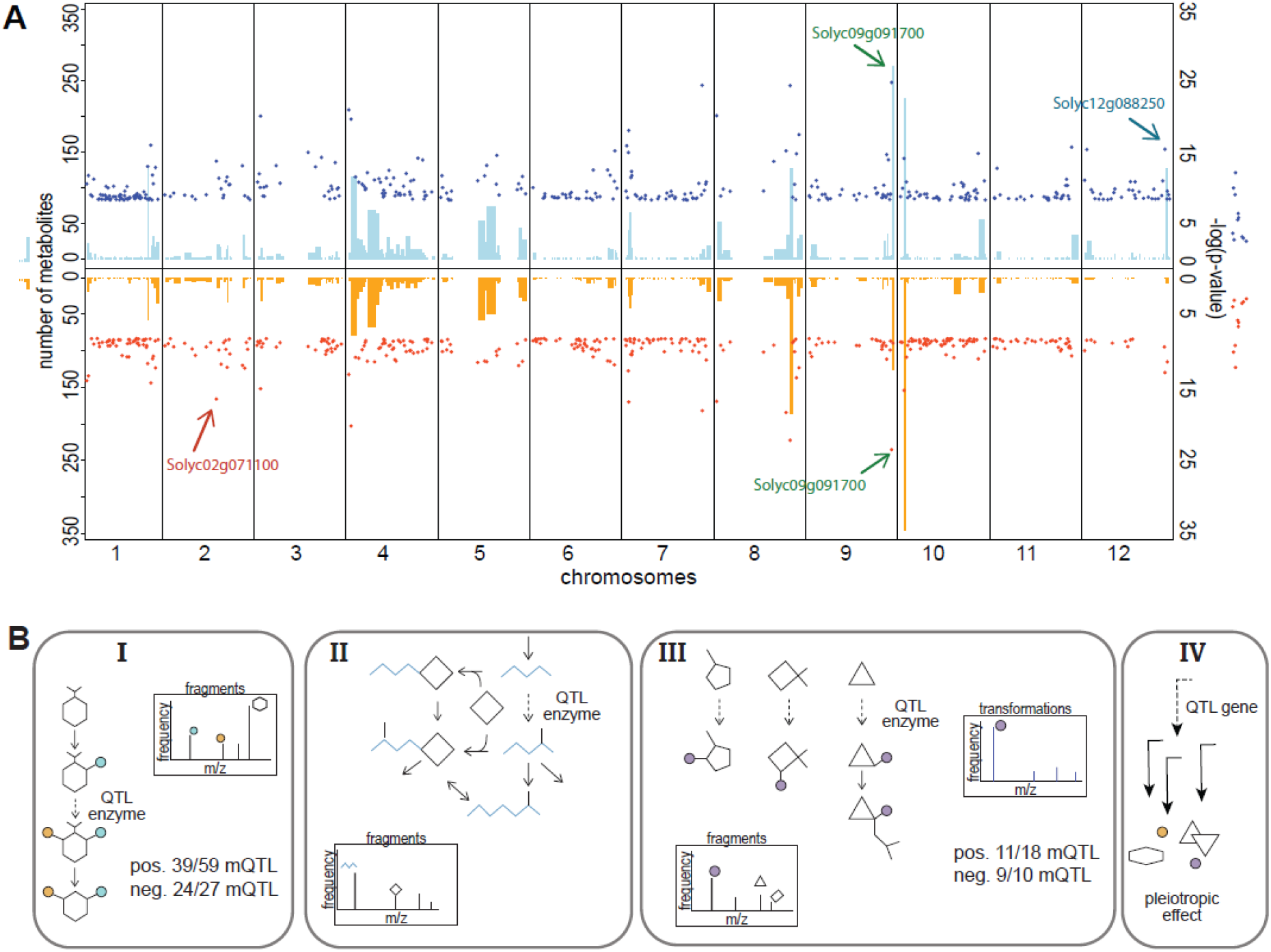
mQTL overview and scenarios. A. Each detected mQTL is represented as a column placed according to physical location on the tomato genome (x axis). The width of the column ranges from the first to the last significant SNP in the mQTL. The height of the column represents the number of metabolite features in the mQTL feature group, scale noted on the left y axis. The dot accompanying each mQTL column represents the significance of the highest SNP-feature association in the mQTL, scale shown on the right y axis. Results of mapping in control conditions are in the top panel in blue, and from water deficit in mirror image in the bottom panel in orange. The mQTL and genes studied in the following sections are marked by arrows. B. Graphical representation of the scenarios discussed in the text for mQTLs with large mQTL feature groups. Structures and colors represent arbitrary molecular substructures. Inset plots are simplified commonality spectra expected from various scenarios. In scenario IV, the pleiotropic effects are not expected to manifest in a typical commonality spectrum, so it is not drawn.

Across all mQTLs, 826 features had repeated genomic associations, often with multiple features co-associating with the same mQTL. Of all features with any significant genomic associations, we were able to obtain ddMS^2^ spectra for 1,445 (55%) in positive, and 750 (48%) in negative ionization mode. However, only 13 features in mQTL feature groups were annotated with good confidence.

Our approach was to be as inclusive as possible, to make maximal use of current high-throughput detection methods. In this way, the results are not biased toward well-studied pathways and may reveal insights into previously uncharacterized compound families. We see value in publishing detailed searchable genomic associations of unannotated features, so that the scientific community can use them as knowledge of compound structures expands. Indeed, we were able to obtain and analyze large amounts of data (Fig. 1A). However, this multitude of data required new approaches to highlight and prioritize relevant information for study. We therefore set out to develop methods for functional characterization of feature groups based on their genomic associations, alongside functional gene characterization based on associated feature groups.

### Commonality analysis: concept and method development

QTLs are usually caused by polymorphisms linked to one causal polymorphism, which primarily influences one causal gene^5,13,15^. Although not always the case, we assumed one causal gene for each mQTL in our working model. When multiple features co-associate with one mQTL, this could arise from various scenarios: I) changes to an enzyme in a pathway which influences upstream and downstream metabolites. II) changes to an enzyme within an interconnected metabolic network. III) changes to an enzyme with multiple substrates. IV) changes to a regulator which indirectly affects metabolism, such as a transcription factor. These various scenarios are expected to produce distinct patterns among the features of the mQTL feature group (Fig. 2B). In detail:

Scenario I) The mQTL-influencing gene is an enzyme which catalyzes a reaction within a defined, linear or weakly branched specialized metabolism pathway. In such a case we expect to see altered abundance of all the compounds in the pathway, which will then be co-associated with the mQTL. For example, a loss-of-function or reduced-activity allele would be associated with an accumulation of precursors and reduction in downstream products. Consecutive reactions in specialized metabolism pathways usually add/remove/alter various parts of the compounds, but the backbone structure is usually preserved and would therefore manifest in the same fragment appearing in the ddMS2 spectra of the features in the mQTL feature group (Fig. 2B I). Identifying these highly shared fragments would define the central substructures and help focus annotation efforts on the basal pathway influenced by the mQTL.

Scenario II) The gene is an enzyme involved in primary or interconnected and redundant metabolic pathways. As in scenario I, altered activity in a specific reaction will ripple and cause coordinated changes in upstream and downstream metabolites. However, redundancy may buffer the allele influence and cause multiple low effect mQTL associations, as often seen in mGWAS of primary metabolites^37,49^. Still, if the polymorphism has a very strong effect, we may see multiple feature-mQTL associations. Due to interconnectivity, different groups of metabolites may be affected. In this scenario shared mass descriptors may be found for features with tight associations to the polymorphism, but we may expect several common descriptors shared by a lower proportion of the features in the mQTL feature group (Fig. 2B II).

Scenario III) The gene is an enzyme which catalyzes a reaction on similar compounds involved in several pathways. In this case we would see changes in compounds of various families. Even promiscuous enzymes usually specialize in a specific type of reaction that is often associated with a specific substructure the enzyme can recognize and bind^50,51^. In this case the recognized substructure may result in a fragment shared by many of the features in the mQTL feature group.

Moreover, the specific reaction would add/subtract a substructure with a specific molecular mass which would manifest in a specific m/z difference between CSPP^29,31^. For example, acetylation or deacetylation (C_2_H_2_O/H) would add or subtract m/z=42.01, respectively. Multiple CSPP features would be co-associated with the mQTL, and the reaction-causing m/z difference would be frequent among all calculated paired m/z differences, named transformations hereafter (Fig. 2B III). A transformation is a calculated mass difference and does not necessarily demonstrate a biochemical conversion.

Scenario IV) If the affected gene is a regulator (for example, a transcription factor) it may affect multiple metabolic pathways, usually ones involved together in a coordinated developmental program or response to changing conditions. They may, however, involve very different reactions and compound structures (Fig. 2B IV). Such regulators may be identified by their central role in gene-gene or gene-metabolite correlation networks but are beyond the scope of this paper.

We hypothesize that mQTL feature groups will have a significantly distinct commonality of descriptors, either fragments, neutral losses (NL) or transformations, compared to random groups of features. Further, these distinct commonly shared descriptors will assist in identifying the mQTL causal gene, reaction and pathway. We set out to test this hypothesis by grouping ddMS^2^ spectra of the features in each mQTL feature group and identifying the frequency of descriptors in the group. We visualized the commonality, the set of descriptor frequencies in a group, as commonality spectra: plots that display, for each m/z value (x axis), the frequency of that descriptor, i.e. the proportion of features in the group that carry it (y axis). Because of the different nature of features and fragments detected in positive or negative ionization mode, the analysis and representation were done separately for each ionization mode. Next, we developed a statistical framework to define the significance of mQTL-based commonality compared to ungrouped features (Supp. Note 1: development of the statistical framework for commonality analysis).

### Descriptor frequencies in mQTL feature groups differ from those in all leaf features

#### Fragments

Prior to studying the commonality of mQTL feature groups, we identified the most common fragments in all collected ddMS^2^ spectra (Supp. Table 5). In positive mode the most common fragments 85.03, 92.20, 127.04, 97.03 were found in 22%, 21%, 18%, 17% of all features with ddMS^2^ spectra, respectively (Fig. 3A). In negative mode the most common fragments 59.01, 101.02, 89.02, 71.01, 113.02 are hexose-derived fragments (Fig. 3B). They were found in over 25% of all features with ddMS^2^ spectra, showing the prevalence of hexosylated (usually glycosylated^52^) metabolites. In addition, the 92.19 fragment of unknown molecular composition is highly common and found in over 25% of spectra. When comparing these overall fragment frequencies to all features influenced by natural variation in our experiment (*i.e.* associated with any mQTL), the pattern was very similar (Supp. Figure 3), indicating the diversity of metabolites and pathways influenced by natural variation. The presence of the most common fragments in such a large proportion of features may cause them to overshadow the mQTL-specific fragments. To focus on mQTL-specific common fragments, these overall common fragments are shown in gray in the following plots and are not counted as most common or outstanding for mQTLs. The frequencies of the top 1,000 fragments are listed in Supp. Table 5.

**Figure 3:**
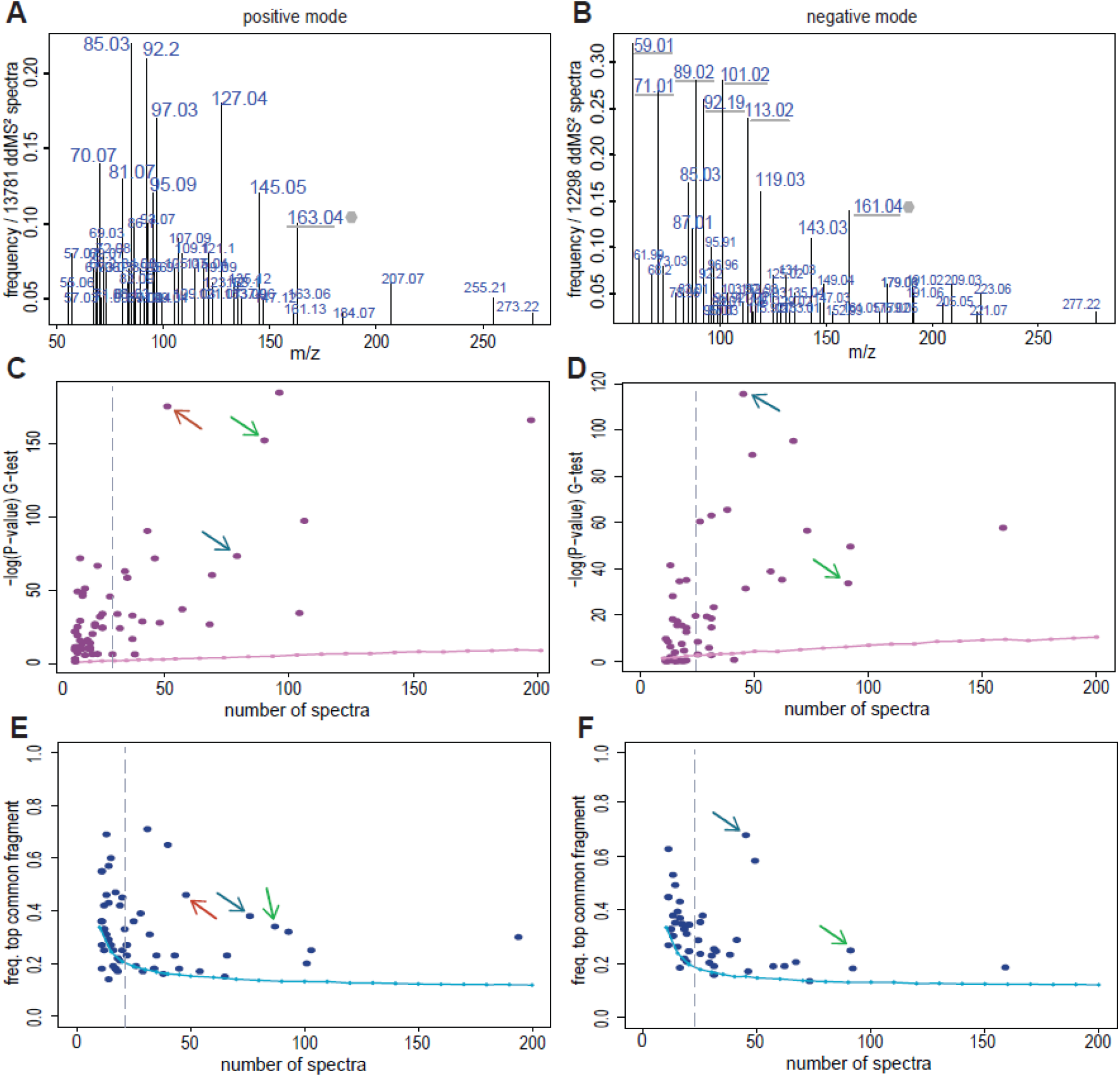
Fragment commonality of mQTL feature groups. A and B. The most common fragments found in all ddMS² spectra obtained from tomato leaf metabolite features detected in positive (A) or in negative (B) ionization mode. The m/z value of each fragment is indicated above the corresponding bar, whose height indicates the frequency of ddMS² spectra which contain the fragment out of all detected ddMS² spectra. Due to the excess of fragments some numbers overlap, the ranked list of fragments and their frequency can be found in Supp. Table 5. Hexose (gray hexagon) derived fragments are marked with gray underline. C and D. The significance level of mQTL fragment commonality indicating how different the commonality of an mQTL feature group is compared to the commonality of all ddMS² spectra. Shown for features detected in positive (C) or in negative (D) ionization mode. The pink line denotes the significance threshold derived from the 5^th^ percentile of random ddMS² spectra sampling with corresponding number of ddMS² spectra. Dashed gray lines note the suggested minimum number of spectra for a confident analysis. E and F. The frequency of the most common fragment of each mQTL feature group (excluding the overall most common fragments from panels A-B), for features detected in positive (E) ionization mode or in negative (F). The light blue line denotes the outstanding threshold, the mean top frequency of random ddMS² spectra sampling with corresponding number of ddMS² spectra. Dashed gray lines note the suggested minimum number of spectra for a confident analysis. Colored arrows indicate the mQTLs studied in the following sections, in accordance with Fig. 2.

We treated the frequencies of fragments in all spectra as the expected values to test whether mQTL feature groups had a significantly distinct commonality (Supp. Note 1). Confident analysis required groups of at least 20 ddMS^2^ spectra. Almost all high-impact mQTLs, those above this size, had a significantly distinct fragment commonality, compared to the α<0.05 significance threshold determined by random permutations (Fig. 3CD), which would match scenarios I or II. Moreover, most high-impact mQTLs (24/27 in positive mode and 19/23 in negative) also had outstanding fragments, whose frequency exceeded the mean top frequency of size-matched random groups (Fig. 3EF), thereby supporting our hypothesis.

#### NL

The fragments detected in ddMS^2^ spectra are only the charged fragments of the molecule. However, the uncharged fragments can be estimated by calculating the NLs within each spectrum^34^. We calculated NLs for each spectrum as the m/z differences between all fragments including the precursor m/z. The resulting set of NLs for each spectrum is its NL pseudospectrum. These NLs can also be treated as descriptors of features, and the most common NL could supplement the fragmentation commonality in highlighting metabolites’ uncharged backbone substructures. NLs have been successfully used to annotate metabolite substructures and identify metabolic genes associated with them^34^. Similarly to fragments, we identified the frequency of NLs in all collected ddMS^2^ spectra and tested the difference in frequency of NLs in mQTL feature groups (Supp. Figure 4; Supp. Table 6).

Because NLs are calculated and not directly measured, NL pseudospectra contain more noise derived from calculated differences which do not actually represent parts of a molecule. For this reason, the statistical power of the commonality analysis is reduced, and requires at least 50 ddMS^2^ spectra (Supp. Note 1) for confident analysis. It is tempting to consider only the annotated NLs (Supp. Table 7), which are likely true molecular substructures. However, one strength of commonality analysis is that it highlights biologically relevant NLs over random artifacts of the calculation. This can be seen in high-impact mQTLs, which had a significantly distinct NL commonality and outstanding NLs compared to random feature groups (8/16; Supp. Figure 4). For example, an NL of unknown structure with m/z of 19.01 is highlighted as an outstanding NL in nine different mQTLs (Supp. Table 4).

#### Transformations

In scenario III, where the mQTL enzyme is involved in multiple pathways, fragments or NL commonality may not be able to highlight a specific backbone. However, the recurring m/z difference, or transformation, between pairs of features in an mQTL feature group may highlight the mQTL-related reaction. Transformations, too, are a calculated descriptor which can have noise due to the “blind” nature of the calculation. Still, the most common transformations found between all pairs of features were known metabolite modifications (Fig. 4AB; Supp. Table 8). For features detected in positive mode, the most common transformations among all measured features are de/hydration, de/hydrogenation and methoxylation (Fig. 4A). In negative mode the most common transformations are methoxylation, de/hydration and acetylation (Fig. 4B). In both modes, these are followed by the calculated transformation of 12.00. However, this does not represent a real biochemical reaction, but rather the many pairs of molecular formulas that differ by one carbon atom. This demonstrates one of the necessary cautions when interpreting the calculated differences. Other common transformations are additions/subtractions of methanol or ethyl groups and sugar (hexose, pentose) modification. The statistical method development (Supp. Note 1) showed that the transformation commonality of mQTL feature groups may confidently be interpreted as significantly distinct from the expected, when applied to groups of at least 50 features (Fig. 4CD). The most common transformation also exceeded the outstanding threshold in most high-impact mQTLs (20/28; Fig. 4EF).

**Figure 4:**
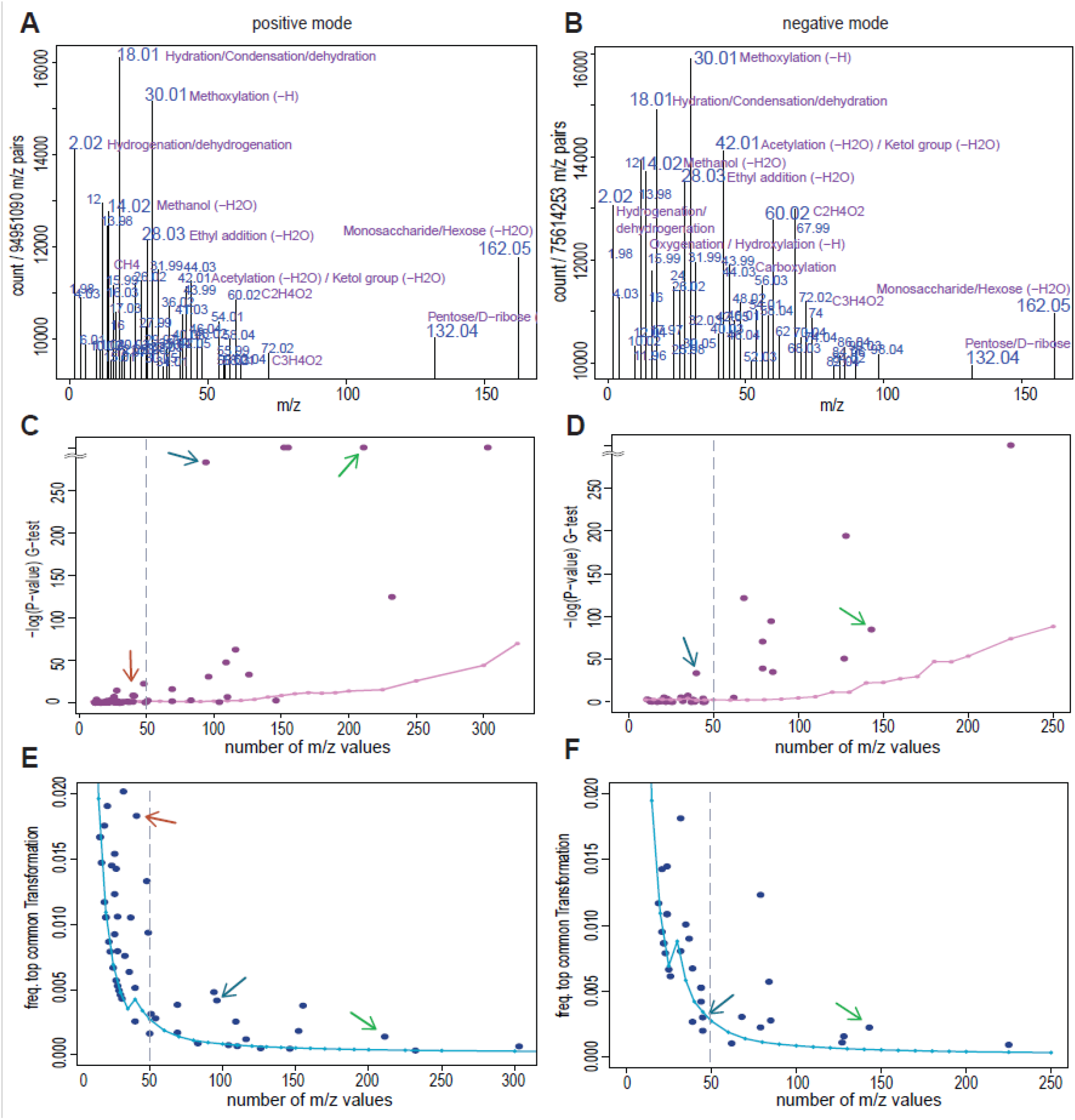
Transformation commonality for identifying mQTLs’ underlying reaction. A and B. The most common transformations (m/z differences) found for pairs of all metabolite features detected in positive (A) or in negative (B) ionization mode. The m/z value of each transformation is indicated above the corresponding bar, whose height indicates the number of occurrences out of all non-redundant feature pairs. Due to the excess of transformations some numbers overlap, the full ranked list of frequent transformations can be found in Supp. Table 8. When available, putative annotations are noted in purple, according to Supp. Table 7. C and D. The significance level of mQTL transformation commonality indicating how different feature pairs of an mQTL feature group are compared to pairs of all features, for features detected in positive (C) or in negative (D) ionization mode. The pink line denotes the significance threshold derived from the 5^th^ percentile of random sampling with corresponding number of feature m/z values. Dashed gray lines note the minimum number of m/z values suggested for a confident analysis. E and F. The frequency of the most common transformation of each mQTL feature group (excluding the overall most common transformations from panels A-B), for features detected in positive (E) or in negative (F) ionization mode. The blue line denotes the outstanding threshold, the mean top frequency of the most common transformation in random m/z samplings, with a corresponding number of feature m/z values. Dashed gray lines note the minimum number of m/z values suggested for a confident analysis. Colored arrows indicate the mQTLs studied in the following sections, in accordance with Fig. 2.

These results support our hypothesis that mQTL feature groups have distinct commonly shared descriptors. In the following sections we demonstrate the utility of this approach for highlighting previously uncharacterized metabolic gene functions.

### Highly common fragments in an mQTL feature group point to a sinapoyl-hexose acyltransferase

An mQTL on chromosome 12 had the most significantly distinct fragment commonality and the most outstanding fragments among features detected in negative mode, and also a significantly distinct commonality and high fragment frequencies in positive mode (blue arrow in Fig. 3C-F). The mQTL was associated with 134 features, mostly in control conditions and very few in WD (Supp. Table 9). We present an in-depth study of this mQTL as an example of how mQTL commonality can reveal novel functions of genes in metabolic pathways.

The mQTL spanned 63,464,146-64,404,847 bp on chromosome 12 (Fig. 5A). The majority of features were most significantly associated with single nucleotide polymorphism (SNP) S12.2059255 at 63,718,397 bp.

**Figure 5:**
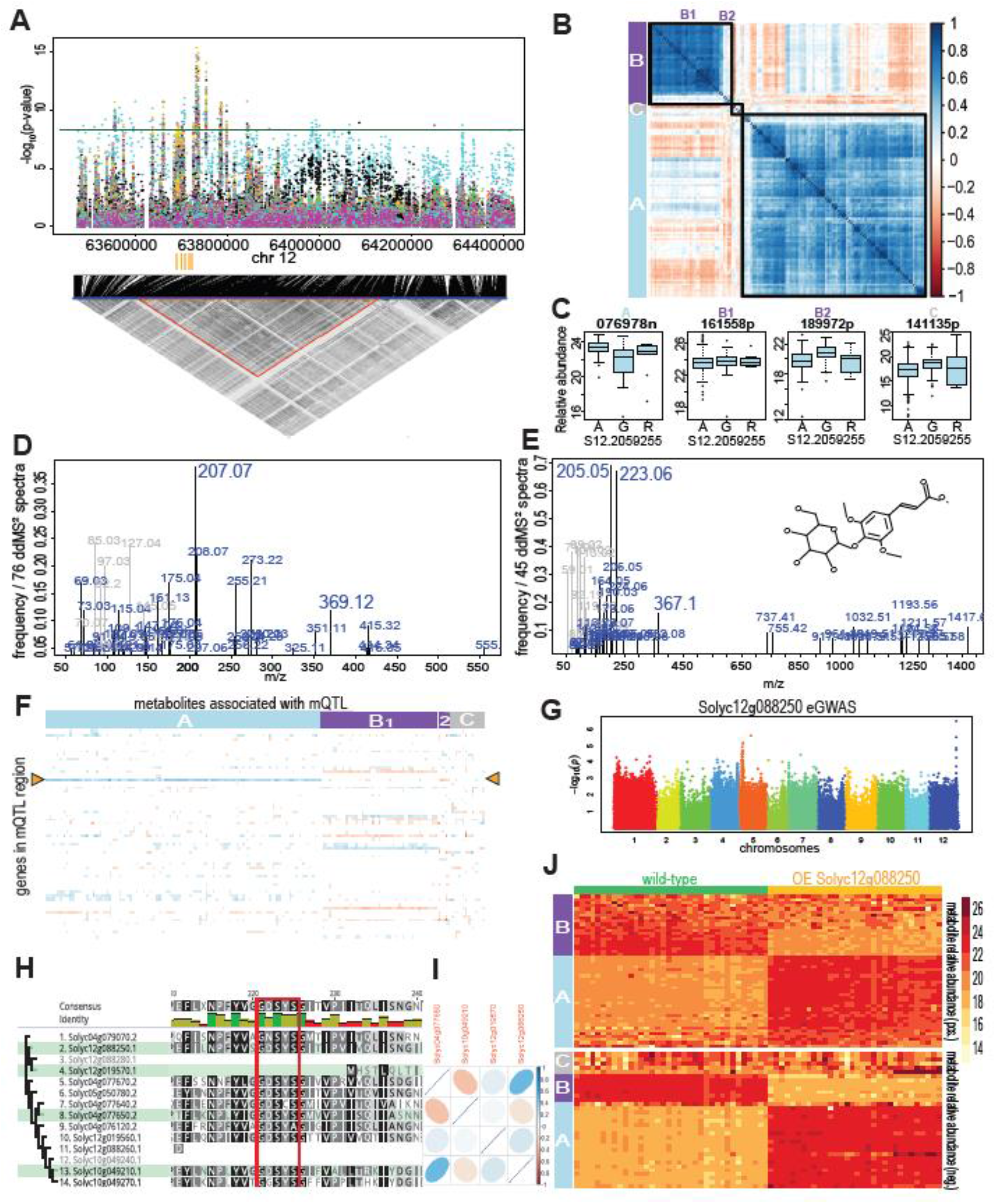
Study of chromosome 12 mQTL based on common fragments. A. A. Manhattan plot of the significant mQTL region in control conditions. Each color represents the mGWAS significance of one of the metabolite features associated with the mQTL. The lower panel shows the linkage disequilibrium analysis of all SNPs in the extended mQTL region. The red triangle indicates the linkage group of the mQTL significant SNPs. Orange rectangles denote the genomic location of serine carboxypeptidase-like (SCPL) family genes. B. Feature-feature correlations for all features of the mQTL feature group in control conditions. Features are ordered and squares are drawn according to the similarity calculated by hierarchical clustering analysis (HCA). Color scale represents the R^2^ coefficient for feature pairs. HCA-based group names were defined according to correlations in control conditions and are noted on the side bar. C. The effect of the most significantly associated SNP, S12.2059255, alleles on relative feature abundance of a representative feature from each HCA group. D and E. Commonality spectra of the chromosome 12 mQTL feature group highlighting the ddMS^2^ fragments detected in positive (D) or in negative (E) ionization mode. The fragments most common among all features are presented in gray. In E, fragments in gray are hexose-derived. Inset in E: molecular structure of sinapoyl-glucose. F. Correlations between expression of genes in the mQTL region (in rows) and features (in columns) significantly associated with the mQTL in control conditions. Color scale represents the R^2^ coefficient, as in panel B. Yellow triangles point to the selected candidate gene, Solyc12g088250. G. Genome wide association (Manhattan plot) of Solyc12g088250 relative expression in control conditions H. Phylogenetic clustering of genes from the SCPL family based on protein sequence alignment. The amino acids of the conserved family motif are marked by a red square. Genes expressed in the tomato leaf samples are highlighted in green. I. Pairwise gene expression correlation of the leaf-expressed genes of the SCPL family. Color scale represents the R^2^ coefficient, as in panel B. J. Heatmap of the changes in features of the mQTL feature group in plants overexpressing Solyc12g088250 (golden top bar) compared to wild-type (green top bar). Features are ordered by HCA groups from panel B.

To better understand the relations between the multiple mapped features and how they may be affected by the mQTL alleles, all features were grouped by pairwise correlation hierarchical cluster analysis (HCA) (Fig. 5B). The first and largest group (designated group A) was mostly negatively correlated to group B. However, many features of group B (group B1) which had mixed correlation to group A also had an mQTL peak with SNP S12.2053749 (position 63,540,503 bp) and may not be directly related to the mapping of other features. Group C had poor correlations with the other groups, and inconsistent mapping patterns which may also be a result of closely located associations mistakenly grouped in the same mQTL. The opposite relations of the HCA groups to each other and to SNP S12.2059255 alleles (Fig. 5C) may suggest they are grouped into the precursors and the downstream products of the mQTL-associated reaction.

To focus annotation efforts on the substructures most central to the pathway affected by the mQTL, we present commonality spectra of the mQTL feature group for features detected in positive (Fig. 5D) or negative ionization mode (Fig. 5E). The outstanding fragments in negative mode were 205.05 and 223.06, present in 68% of mapped features, and multiple hexose fragments (Fig. 5E, in gray). Several of the most common fragments in positive and negative could be paired by the typical mass difference of 2 between [M+H]+1 and [M-H]-1 ions. Consistently, fragment 207.07 was the most common in positive mode, present in 38% of features.

Based on experimental^47,53^ and predicted fragmentation^54^, the fragment pairs 205.05/207.07 and 223.06/225.05 are typical of a sinapoyl substructure. The expected 225.05 had low abundance in our data. The fragments 367.10 and 369.11, also among the most common fragments in negative and positive modes, respectively, match sinapoyl-hexose (Fig. 5E, inset). The presence of hexose (162.05) in many of the features in the mQTL feature group is also observable in the NL commonality in both positive and negative modes (Supp. Figure 5AB), as is the common presence of the sinapoyl (206.06, 224.07) and sinapoyl-hexose (368.11) in negative spectra NLs (Supp. Figure 5B).

Of the features associated with the mQTL, only one was directly annotated: it matched 1,2-di-O-sinapoyl-beta-D-glucose in the GNPS library (Supp. Table 9), and four others were linked to sinapoyl-hexose in the FBMN (Supp. Table 10). However, we were able to identify the typical fragment in 58 of the features in the group (of the 97 with available ddMS2 spectra) and could therefore extend the substructure annotation to 43% of the mQTL feature group.

The mQTL region included 28 genes, but none was annotated to catalyze an expected metabolic reaction. Correlations between features of the mQTL feature group and relative expression of genes within the mQTL region pointed to a gene, Solyc12g088250, with many significant correlations (p-value < 10-7, |r|>0.4; Fig. 5F). Solyc12g088250 is located at 63,710,096 bp, right at the peak of the mQTL and is annotated (ITAG2.4) as a serine carboxypeptidase, one of four tandem genes with the same annotation in the mQTL region. Analysis of gene expression levels by GWAS revealed a cis expression QTL (eQTL) co-locating with the mQTL (Fig. 5G), suggesting that a polymorphism affecting Solyc12g088250 expression underlies the mQTL.

We searched in Plant Metabolics Network (http://www.plantcyc.org) databases and the literature^55–57^ for enzymes involved in sinapoyl-hexose metabolism and found multiple mentions of enzymes of the serine carboxypeptidase-like (SCPL) family. We therefore performed a detailed homolog phylogenetic analysis (Supp. Figure 5C) which revealed that Solyc12g088250 is in fact part of the SCPL subfamily, a group of metabolic genes with a conserved motif of GD<u>S</u>YS (<u>S</u> is the essential catalytic serine)^56^, with no peptidase activity. The redefined gene family annotation and common substructure annotation supported the involvement of Solyc12g088250 in sinapoyl-hexose transferase activity. We therefore set out to validate the gene’s activity by overexpressing (OE) Solyc12g088250 in tomato plants.

#### Validation of gene-metabolite association by Solyc12g088250 overexpression

The Solyc12g088250 OE plants of five independent transformation lines (Supp. Figure 6A) were visually comparable to the wild-type (WT) plants (accession Ailsa Craig), but metabolic analysis revealed clear differences (Supp. Figure 5D). Of the 134 features associated with the mQTL, 59 features were also found in the OE experiment. Most of them (43/59) were significantly altered in OE samples compared to WT (Fig. 5J).

Features of the large HCA group A were significantly increased by OE, in agreement with the positive correlation between gene expression and their feature abundance seen in the natural population. The features in group B2 were strongly associated with the mQTL lead SNP, and the four which were found in the OE experiment displayed decreased abundance in OE samples, as expected from their negative correlation to group A and to Solyc12g088250 gene expression.

Additional features which were not associated with the mQTL were affected by the OE. Partial least squares discriminant analysis (PLS-DA) was used to identify which features contributed most to the difference between OE and WT samples. We selected the top 1% of contributing features (168 in positive and 85 in negative ionization modes) (Supp. Figure 5E). These were also significantly different (t-test) between OE and WT. Commonality analysis of these features revealed hexose fragments and sinapoyl-hexose fragments (205.05, 223.05 and 367.1) among the most common in negative mode and sinapoyl (207.07) in positive mode (Supp. Figure 5F). However, in addition to the expected fragments, reproducing the central mQTL substructures, additional fragments were common among the features most distinguished between OE and WT. Possibly, strong OE reveals a more extended effect of the gene on metabolites whose abundance changes in the natural variation are mitigated by interconnected metabolic networks. Alternatively, OE of a gene may at times produce unnatural promiscuous activity, which extends beyond the naturally occurring reactions^58,59^. Overall, the OE experiment confirmed that the SCPL gene Solyc12g088250 affects the features of the mQTL feature group within the sinapoyl-hexose pathway.

### Shared and divergent activity of two SCPL genes

Twelve genes in the tomato genome fell within the SCPL subfamily clade of the phylogenetic tree (Fig. 5H). Four of them were expressed in leaf samples in this experiment (Fig. 5H, green highlight). Solyc12g019570 encodes a truncated protein without the GD<u>S</u>YS motif and is probably a pseudogene. Solyc10g049210 is an ortholog of Sopen10g020280 from tomato relative *S. pennellii*, previously characterized as an isobutyrylglucose:isobutyrylglucose acyltransferase (GAT)^60^.

Interestingly, the expression of Solyc04g077650 was positively correlated with that of Solyc12g088250 (Fig. 5I). The sequence similarity and coordinated expression raised the question of whether these genes were redundant or served distinct metabolic reactions. We hypothesized that commonality analysis could be utilized to highlight important descriptors for features grouped by methods other than co-association. Here, we used it in a reverse-genetics approach to identify overlap and specificity between related SCPL enzymes.

To this end, we produced plants overexpressing Solyc04g077650 (Supp. Figure 6B). Again, PLS-DA was used to identify which features contributed most to the difference between OE and WT samples (Fig. 6A). The fragment commonality of this OE-affected feature group was significantly (p-value < 10^-16^) distinct from the expected. The most common fragments were similar to the ones found in commonality of the Solyc12g088250 OE, albeit different in order of frequency. To further refine the way commonality analysis helps us understand feature-gene relations, we divided the group of OE-affected features into subgroups: features increased by OE, expected to be downstream products of the enzyme reaction, and those decreased by OE, expected to be upstream precursors. Side-by-side comparison of the commonality of predicted precursors and products of the two genes is presented in Fig. 6.

**Figure 6:**
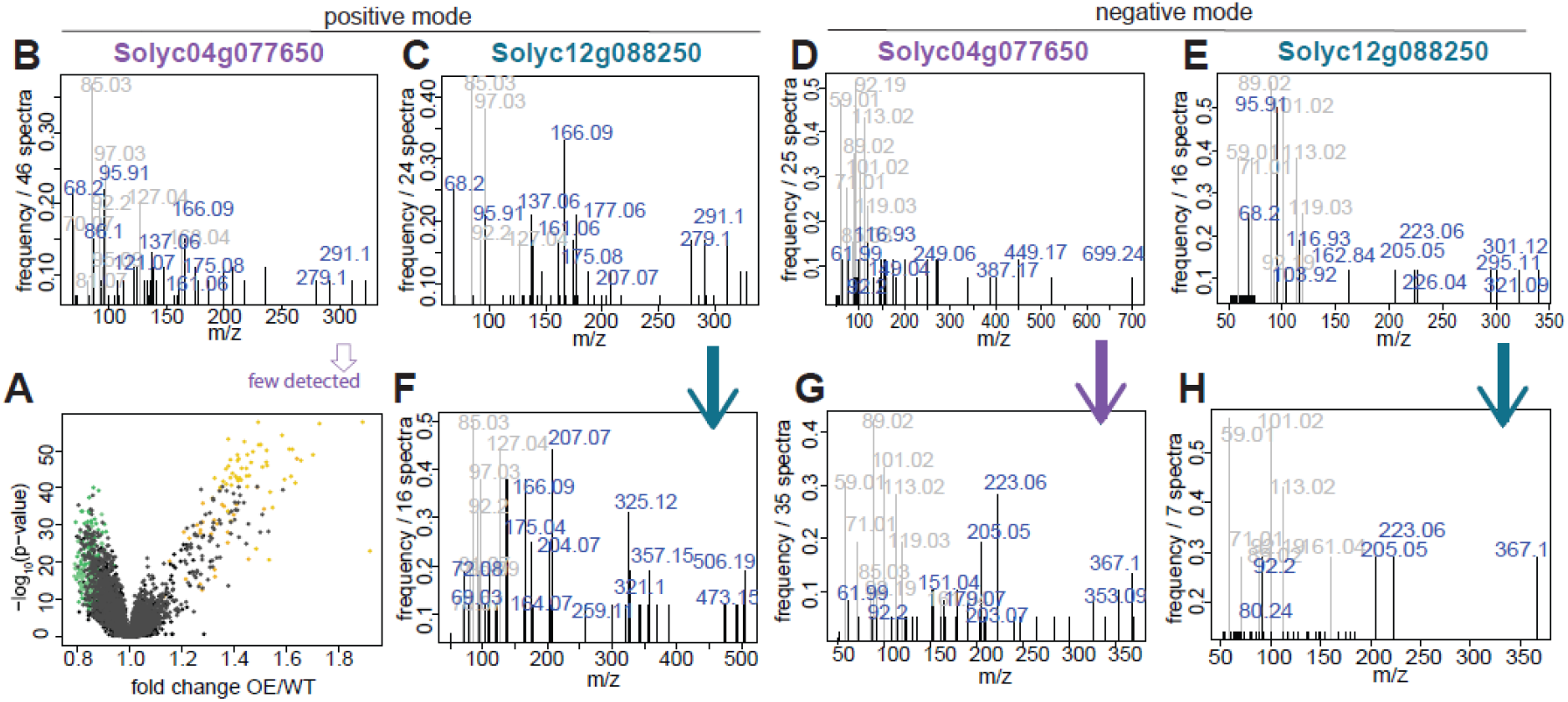
Fragment commonality of predicted precursors and products of two serine carboxypeptidase-like genes. A. A. Volcano plot displaying the significance of a t-test comparing metabolite feature abundance between plants overexpressing (OE) Solyc04g077650 and wild-type (WT; Ailsa Craig accession) against the fold change (after log2 transformation) of the same comparison. Features which contributed most (top 1%) to the separation of OE and WT in PLS-DA are colored golden (increased in OE) or green (decreased in OE). B-E. Commonality spectra of fragments of features decreased by OE of Solyc04g077650 (B, D) or Solyc12g088250 (C, E) in positive (B, C) or negative (D, E) ionization modes. F-H. Commonality spectra of fragments of features increased by OE, expected to be the products of the reaction as denoted by the arrows, of Solyc04g077650 (G) or Solyc12g088250 (F, H) in positive (F) or negative (G, H) ionization modes.

For both genes many precursors and products contain hexose fragments. The typical sinapoyl-hexose fragments are found in the products of both genes but in the precursors of Solyc12g088250 only, suggesting that this enzyme may add sinapoyl and hexose to molecules already containing these groups (Supp. Figure 5G). In addition, apart from hexose fragments, the fragments found in more than 10% of precursors are different between the genes, suggesting they perform a similar function of sinapoyl acyltransferase with different substrate specificity.

Overall, the most common fragments point to sinapoyl-hexose as a core substructure in the pathway altered by the mQTL. Correlation and co-locating eQTL of Solyc12g088250 gene expression suggest that genomic variation alters expression, in turn influencing enzyme activity levels and metabolite abundance in the sinapoyl-hexose pathway. Solyc12g088250 was misannotated by automated annotation. Without the rapid identification of the sinapoyl-hexose substructure by commonality analysis, this connection would have been harder to make. We further demonstrate the utility of commonality analysis to quickly highlight metabolic pathways affected by experimental genetic modifications, which may be used in forward and reverse-genetics approaches to identify metabolic gene functions.

### WD-specific regulation of a putative transporter influences accumulation of multiple nitrogen-containing metabolites

The next mQTL had one of the most significantly distinct fragment commonalities in positive ionization mode, but its interpretation was more challenging since the main fragment structure is unknown. The mQTL spanned 40,443,154-40,914,962 bp on chromosome 2. It was significantly associated with 49 features (Fig. 7AB). In WD, 35 features were associated, five of which were also significantly associated in control, but with lower significance (Supp. Table 11; Supp. Figure 7). In contrast, 14 features were significantly associated only in control (Fig. 7A) but showed the highest association with a SNP 60 kb away from the WD-associated features.

**Figure 7:**
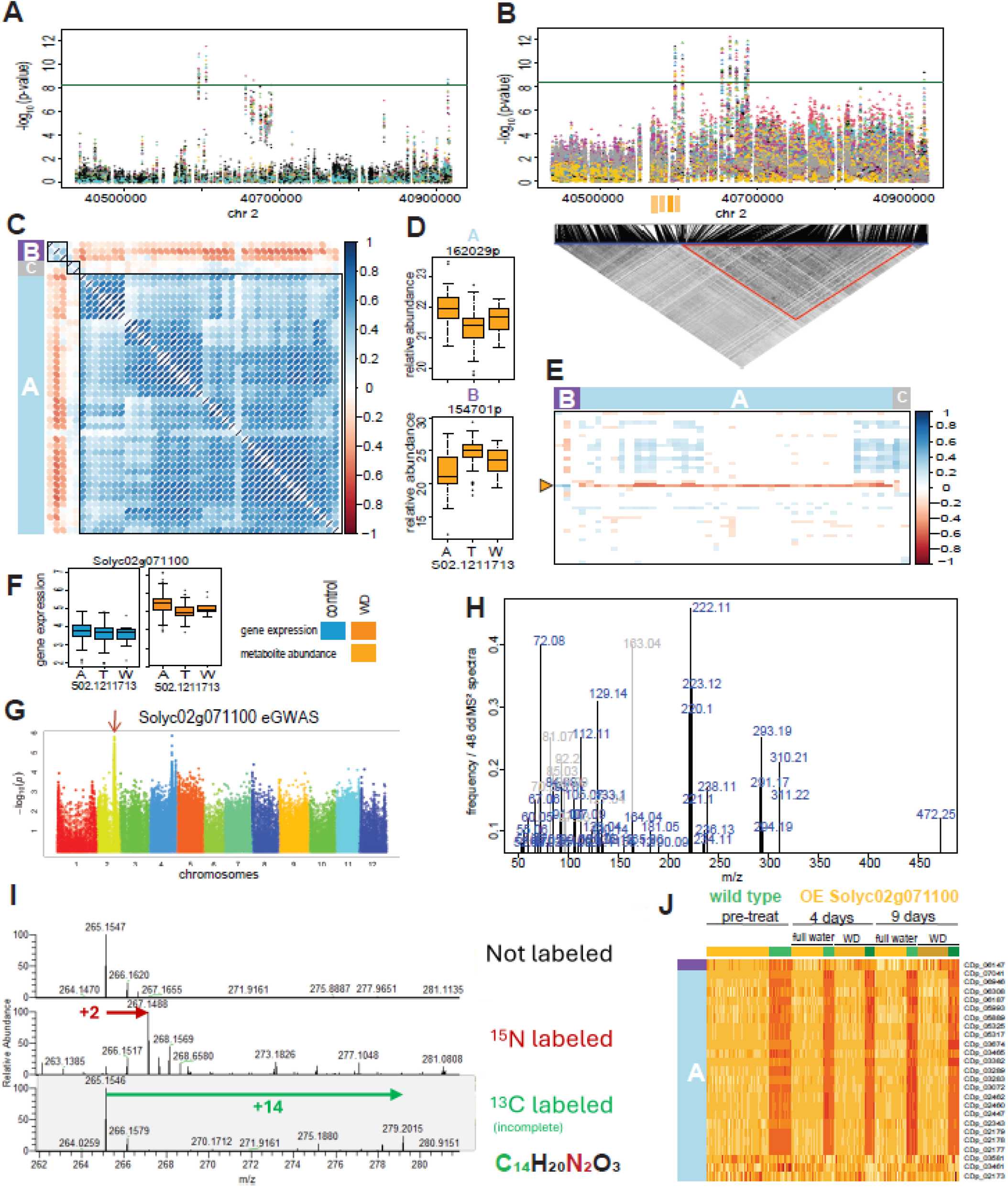
Study of chromosome 2 mQTL and candidate gene A-B. Manhattan plot of the significant mQTL region in control. (A) and water deficit (B) conditions. Each color represents the GWAS results of one of the metabolite features associated with the mQTL. The lower panel in B shows the linkage disequilibrium of all SNPs in the extended mQTL region. The red triangle indicates the linkage group which includes the mQTL significant SNPs. Orange rectangles denote the genomic location of purine permease family genes. C. Feature-feature correlations for all features of the mQTL feature group in water deficit conditions. Features are ordered and grouping squares are drawn according to the similarity calculated by hierarchical clustering analysis (HCA). Color scale represents the R^2^ coefficient for feature pairs. HCA-based group names are noted on the side bar. D. The effect of the most significantly associated SNP, S02.1211713, alleles on relative feature abundance of representative features from the HCA groups in water deficit. E. Correlations between expression of genes in mQTL region (in rows) and features (in columns) with significant association to the mQTL in water deficit conditions. Color scale represents the R^2^ coefficient, when significant. The yellow triangle points to the selected candidate gene, Solyc02g071100. F. The effect of the lead SNP S02.1211713 and of the environment on Solyc02g071100 gene expression. G. Genome wide association (Manhattan plot) of Solyc02g071100 relative expression in water deficit conditions. H. Fragment commonality spectrum of the mQTL feature group, for features detected in positive ionization mode. I. Spectra at retention time 4.44 min derived from ^15^N and ^13^C isotope labeling experiments. The comparison of spectra of this representative feature shows the increase in m/z corresponding to the number of labeled atoms, validating the predicted (C_14_H_20_N_2_O_3_) molecular formula of feruloylputrescine. J. Heatmap of the changes in features of the mQTL feature group in plants overexpressing Solyc02g071100 (golden top bars) compared to wild-type (green top bars), under fully watered or water deficit conditions at the indicated time points. Feature side colors indicate the HCA groups from panel C.

The 49 features of the mQTL feature group were categorized into three groups by correlation-based HCA (Fig. 7C). Groups A and B were negatively correlated to each other and showed opposite effects of the most significantly associated SNP S02.1211713 (Fig. 7D).

Of the 95 genes in the mQTL region, 34 were expressed in the leaf samples. One gene, Solyc02g071100, was highly correlated (p-value < 10^-7^, |r|>0.4) with most features that were associated with the mQTL in WD conditions (Fig. 7E). The expression of Solyc02g071100 across all samples was elevated in WD samples compared to the control (Fig. 7F). The alleles of the lead SNP of the mQTL had a significant effect on the gene expression in WD, but not in control samples (Fig. 7F). This association was further supported by an eQTL for Solyc02g071100 expression co-locating with the mQTL exclusively in WD conditions (Fig. 7G).

Solyc02g071100 is annotated as a purine permease gene with a GO term of purine nucleobase transmembrane transporter activity. It was found to be highly similar to nine other genes in the tomato genome annotated as purine permeases, three of them within the same mQTL region (Fig. 7B). Neither adenine nor guanine (Supp. Table 3) was associated with the mQTL. In fact, all associated features had much higher molecular weight, pointing to more complex compounds.

The majority of associated features were detected in positive ionization mode, so commonality analysis was only possible for these features. The most common fragments in the mQTL feature group were 222.11, 72.08, 163.04, 223.12 and 129.14 (Fig. 7H), but the substructure they are derived from is not known. Among the common fragments of the mQTL feature group many had even nominal m/z values, suggesting nitrogen-containing compounds (nitrogen rule for protonated ions)^61^. The only annotated feature in the group was feruloylputrescine, and two others had some similarity to this compound. Together with the putative annotation of Solyc02g071100 as a purine permease and the association with feruloylputrescine, this led us to hypothesize that the mQTL is involved in the metabolism of nitrogen-containing compounds.

#### Annotation support by stable isotope labeling

To test this hypothesis and improve annotation, we conducted an isotope labeling experiment using ^15^N-labeled plants and ^13^C-labeled calli. The feruloylputrescine annotation, with the C_14_H_20_N_2_O_3_ predicted molecular formula, was confirmed by a shift of the precursor ion m/z of 265.1547 by 1.9941 in ^15^N-labeled samples and by 14.0469 in ^13^C labeled samples (Fig. 7I). We continued investigating all features of the mQTL feature group which were found in the labeled samples in a similar fashion (Supp. Figure 8). Because specialized metabolite production depends on environmental conditions, it is expected to differ in tissue culture. Of the 41 features of the group in positive mode, 20 were found in the ^15^N-labeled samples, and fewer in the ^13^C-labeled calli. All detected features were demonstrated to contain between one and four nitrogen atoms in their molecular formula (Supp. Figure 8).

Of the features in the group, 22 contained the 222.11 fragment. Of these, nine were found in the ^15^N-labeled samples and were confirmed to contain 2-4 nitrogen atoms. We conclude that the mQTL influences nitrogen metabolism, and next validated the involvement of the candidate gene.

#### Validation of nitrogen metabolism function by Solyc02g071100 overexpression

We next set out to validate the role of Solyc02g071100 as the gene underlying the mQTL by OE (Supp. Figure 6D). Since the effect of the mQTL was more pronounced in WD conditions, the OE lines and control WT were subjected to reduced water availability prior to metabolic profiling. As expected, the developmental stage, sampling time and water availability influenced the general metabolic profiles (Supp. Figure 7C). Remarkably, the overexpression of Solyc02g071100 also had a substantial effect on the general leaf metabolic profile (Supp. Figure 7C). Of the 41 features of the mQTL feature group detected in positive mode, 25 were found with high confidence in the OE experiment samples. All but three displayed a strong reduction in OE samples (Fig. 7J), as expected from the negative correlation between gene expression and feature abundance in the GWAS population. The effect of OE was high regardless of the watering regime, probably due to constitutive high expression by the 35S promoter overriding any condition-dependent gene regulation.

Following the strong effect of Solyc02g071100 OE on the metabolic profile, we identified the top 1% of features most influencing the separation between WT and OE using PLS-DA (Supp. Figure 7F). Again, the most common nominal m/z values in this group of features were even, suggesting nitrogen-containing compounds (Supp. Figure 7G). Fragment 222.11 was highly common in OE-decreased features, as in features negatively correlated with gene expression in the natural population, but absent from features increased by OE.

Interestingly, many features which increased in OE had very high m/z values (800-1486), while features with lower abundance in OE had mostly m/z values under 600, indicating that the consequence of this gene’s activity may involve synthesis of more complex compounds rather than degradation. Also, the high m/z values of the most common transformations between features increased versus decreased by OE point to the addition of high-molecular-weight groups to the compounds (Supp. Figure 7H). In support, the only three features in HCA group A (positively correlated with gene expression) have the three highest m/z values among the features of the mQTL feature group (Supp. Table 11).

Taken together, we propose Solyc02g071100 as a WD-induced transporter of nitrogen-containing compounds and hypothesize that it transports the compounds into a compartment where they are metabolized to form high-molecular-weight compounds.

### Outstanding transformation in an mQTL feature group points to a double bond reductase

We next tested the ability of transformation commonality to reveal the reaction underlying an mQTL. One of the mQTLs with the highest frequency of an outstanding transformation in positive and negative modes was found on chromosome 9. It was consistently associated with a high number of features, 271 and 126 in control and WD conditions, respectively (Supp. Table 12).

The mQTL spanned 70,800,000-71,600,000 bp (Fig. 8AB). Detailed examination revealed a strong association with a single SNP in the mQTL located at 70,943,627 bp, with very low linkage (LD<0.1) to nearby SNPs. The SNP was positioned on the last base of the predicted stop codon of Solyc09g091700 (Fig. 8D), eliminating the predicted stop codon (ITAG2.4) and adding four amino acids, Trp-Lys-Lys-Lys, to the end of the protein. Expression levels differed substantially in the population. However, expression was not associated with the lead SNP alleles or correlated to the abundance of features in the mQTL feature group, suggesting the coding polymorphism, not expression differences, causes this mQTL. The reference allele A (genome version SL2.50) was ubiquitous in *S. pimpinellifolium* and *S. lycopersicum* var. cerasiforme, but allele G appeared among heirloom, transitional and modern accessions at a rate of 0.36, 0.18 and 0.27, respectively. The position of the stop codon in the A allele also matches that of other homologous proteins (Supp. Figure 9A) and is likely the more active version.

**Figure 8:**
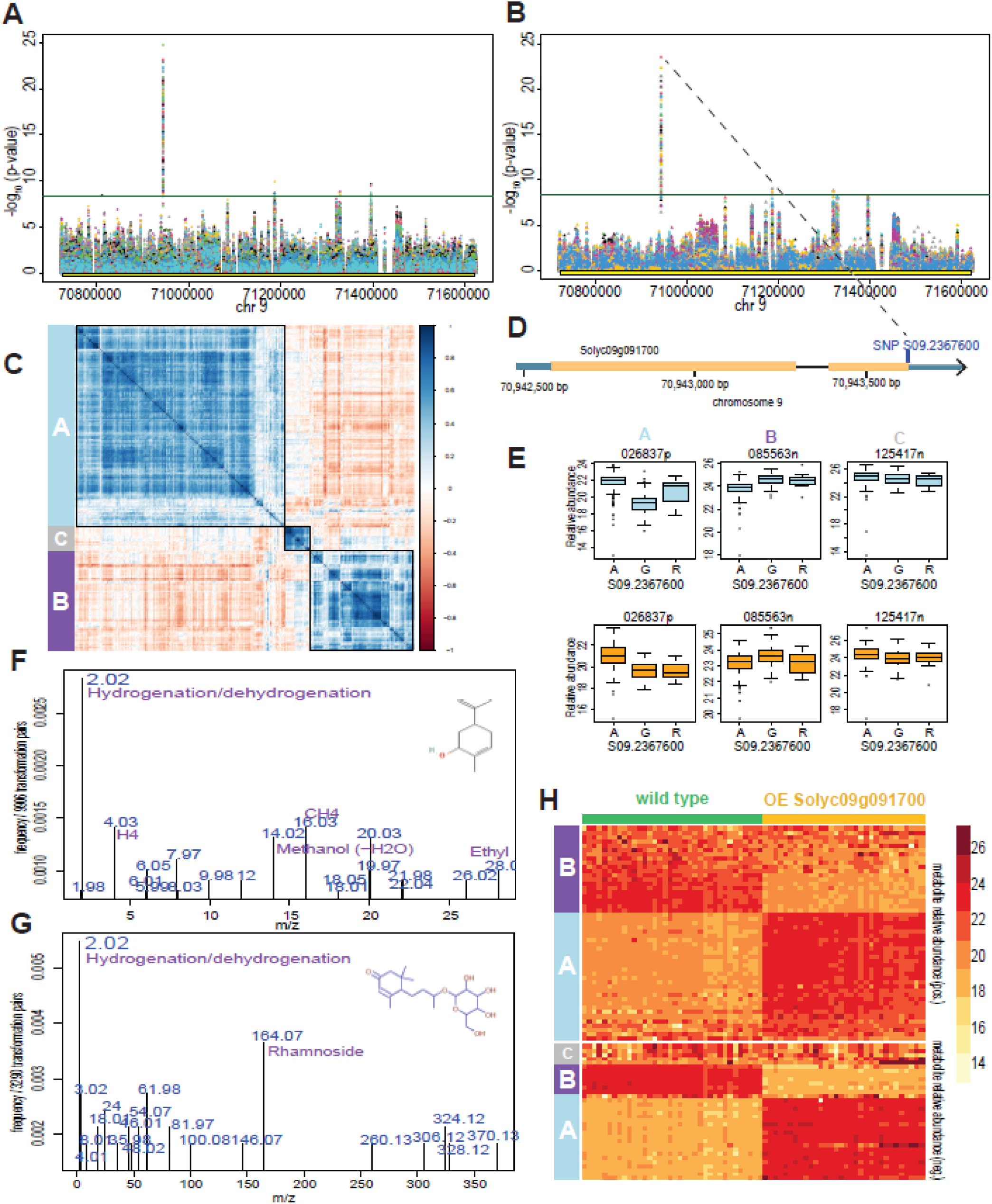
Study of chromosome 9 mQTL based on high transformation commonality. A and B. Manhattan plot of the significant mQTL region in control (A) and water deficit (B) conditions. Each color represents the mGWAS significance of one of the metabolite features associated with the mQTL. C. Feature-feature correlations for all features of the mQTL feature group in water deficit conditions. Features are ordered and squares are drawn according to the similarity calculated by hierarchical clustering analysis (HCA). Color scale represents the R^2^ coefficient for feature pairs. HCA-based group names were defined according to correlations in control conditions and are noted on the side bar. D. Focus on the genomic location of the single highly associated SNP, within the context of the gene Solyc09g091700. E. The effect of the highly associated SNP, S09.2367600, alleles on relative feature abundance of representative features from each HCA group in control (light blue) and water deficit (orange). F and G. Commonality of transformations between features in HCA groups A and B, which are negatively correlated to each other (panel C), for features detected in positive (F) or in negative (G) ionization mode. Inset: molecular structures of annotated metabolites with the typical double-bond of the mQTL feature group in positive (F) or in negative (G) ionization mode. H. Heatmap of the changes in features of the mQTL feature group in plants overexpressing Solyc09g091700 (golden top bar) compared to wild-type (green top bar). Features are ordered by HCA groups from panel C.

The correlation-based HCA of features associated with this mQTL showed two major groups, A and B, negatively correlated to each other, and a third group C, with mixed relations to the first two groups (Fig. 8C). Groups A and B had consistently opposite association with the lead SNP alleles in both conditions (Fig. 8E). Because group A features increase with the likely active allele A, they are probably the products of the reaction, and group B features are likely precursors.

The transformation commonality revealed 2.02 as the most frequent transformation among all feature pairs of the mQTL feature group (Supp. Figure 9BC). Although 2.02 is one of the common transformations found among all features (Fig. 4AB), it stands out with an exceptionally high frequency among features associated with this mQTL. To further refine the transformations most relevant for the enzymatic reaction driven by the mQTL, we calculated transformations between the likely precursors and products. We calculated the commonality of m/z differences between HCA group A and group B features. In this directional analysis, 2.02 stood out at an even higher frequency in both positive and negative modes (Fig. 8FG), suggesting that the mQTL-altered reaction adds 2.02 Da to the precursor compounds. The 2.02 Da m/z difference is consistent with that of 2 hydrogen atoms, or H_2_. Once we identified 2.02 as the transformation central to the mQTL, we also noticed the same mass difference of 2.02 between common fragments of the mQTL feature group (Supp. Figure 9DE).

Based on spectral matches to databases we were able to annotate seven features of the mQTL feature group (Supp. Table 12). An additional seven features had partial or substructure matches, three of which included a carveol substructure. The carveol main library fragments were among the fifty most common fragments of the mQTL feature group: 135.11, 119.085 and 137.096 (Supp. Figure 9D), and at least one was present in 46 (56%) of the features that had a ddMS^2^ spectrum in positive ionization mode. All structures we annotated for features of the mQTL feature group include a cyclohexene or cyclohexane ring (Supp. Figure 9J).

The gene affected by the lead SNP, Solyc09g091700, is annotated as an alcohol dehydrogenase, zinc-containing gene, with two specific GO terms: 2-alkenal reductase [NAD(P)] activity and alcohol dehydrogenase (ADH) [NAD] activity. Solyc09g091700 was putatively assigned to ten alcohol dehydrogenase (EC 1.1.1.1) reactions in the SolCyc database^62^, producing aldehydes. However, none of the annotated features in the group included an aldehyde. The closest Arabidopsis homolog, AT5G16970 (top BLASTP hit in TAIR, 49% identity, 69% positives), is annotated as a 2-alkenal reductase (EC 1.3.1.102). A similar gene, *Nicotiana tabacum* double bond reductase (*NtDBR*), has been shown to catalyze reduction of double bonds on a variety of substrates^63^, with similarity to the structures we found (Supp. Figure 9J). These reactions incorporate H_2_ into the molecule, consistent with the common 2.02 transformation of the mQTL.

#### Validation of Solyc09g091700 activity by overexpression

Next, we set out to validate this hypothesis by OE of Solyc09g091700, using the A allele coding sequence (Supp. Figure 6C). We found very high similarity between commonality spectra of the mQTL feature group and of the top 1% of features contributing most to the separation between WT and OE samples (Supp. Figure 9D-I). This allowed a quick confirmation that Solyc09g091700 affects the features associated with the mQTL. Of the 355 features which were associated with the mQTL, 83 were found again in the OE experiment. The majority of them (70/83) were significantly different between OE plants and WT (accession Micro-Tom) (Fig. 8H). Features in HCA group A significantly increased in OE, while those in group B decreased. This agrees with A being the functional allele, with group A features as products and group B features as precursors. Directional transformation analysis between features increased by OE and those decreased highlighted 2.02 as the most frequent transformation, as in the directional transformation analysis of the mQTL.

Taken together, we propose that Solyc09g091700 functions as a double bond reductase, highlighted by the 2.02 transformation. It likely accepts multiple substrates, which often include cyclohexene or cyclohexane substructures. The coding variation in this gene alters enzyme activity and influences the abundance of hundreds of features in both WD and control conditions.

## Discussion

### Application and limitations of commonality analysis

In this study, we set out to reveal novel metabolic regulation in tomato leaves using natural variation and environmental stress. The combination of genomic, transcriptomic and metabolomic variation in natural populations has been successfully used in recent studies to identify metabolic gene functions^8^. However, studies have often been limited to annotated or partially annotated compounds. Some previous studies have identified and quantified thousands of features but proceeded to map and study only hundreds of them^36,37,64^. Currently, new discoveries are less limited by data acquisition and more by processing and prioritizing deeper investigation. In this era of big data, the commonality analysis presented here can facilitate gene-metabolite association and functional annotation of previously understudied metabolites.

It has been shown that substructure annotation or semi-annotation is useful for discovery of novel gene-metabolite associations^34^. Zhu *et al.* studied genomic hotspots where multiple metabolites sharing the same identified substructure, a fragment or NL, co-associate. The FBMN approach groups features by fragment similarity to identify the underlying structures and group related metabolites. However, when grouping based solely on similarity, not all pathway metabolites which share the same fragments will be tied together in the FBMN, as seen by the dispersed placing of sinapoyl containing features linked to the Solyc12g088250 mQTL. The commonality analysis combines the advantages of both approaches, grouping by genomic co-association and using the outstanding fragments to define the pathway backbone structure. Grouping by genomic or other biological association may provide a more direct route to biological interpretation.

We show that most high-impact mQTLs have a significantly distinct fragment commonality and outstanding fragments suggested to belong to the backbone of affected pathways. Still, commonality analysis is a prioritization and interpretation tool, not a replacement for metabolite identification or functional validation. The simplicity of the proposed calculations, however, allows for quick evaluation of fragment, NL and transformation commonality for each mQTL or other studied group. Nevertheless, the method is not universally effective. Depending on the genetic polymorphism and the structure of the metabolic pathway, some mQTLs may have no outstanding descriptor.

The major limitation of the method is that it requires a large mQTL feature group with ddMS^2^ spectra for its features. Another is that the statistical framework does not currently correct for the testing of multiple mQTLs (or other groups) in the experiment.

The use of NL commonality comes as a complementary method to using fragments for semi-annotation. However, the presence of noise from calculated differences which are not substructures can hamper the analysis. It therefore requires an even higher number of ddMS^2^ spectra for a meaningful analysis. If available, the commonality method has the advantage of identifying highly recurring, and therefore possibly important, NLs, even if they are not annotated. For example, nine mQTLs have an outstanding NL of 19.01 in 37-61% of the features in the group (Supp. Table 4), with an unknown formula, but perhaps worth investigating.

The transformation commonality analysis builds upon the CSPP approach^29,31^, where the m/z difference between two features points to the biochemical reaction turning one to the other. However, instead of suggesting a reaction for each pair of features, it draws upon the dominant transformation for the whole group, or between groups, to suggest the reaction influencing them all. Transformation analysis benefits from using precursor ion m/z values and therefore does not require ddMS^2^ spectra. However, it shares the disadvantage of using calculated descriptors which do not all represent true metabolic differences.

The poorly correlated subgroups of features observed at several loci illustrate both the risk of automated grouping by co-association and the ability of dominant common descriptors to remain detectable despite some unrelated features.

The highlighted descriptors can then be evaluated in combination with other available information, such as gene expression correlations, gene annotations and similarity to annotated compounds to identify targets for investigation.

### Common sinapoyl fragments reveal an SCPL-associated metabolic pathway

A clear example can be found with the mQTL on chromosome 12. The two outstanding fragments quickly pointed to the sinapoyl-hexose pathway. The gene expression correlation implicated Solyc12g088250, which we identified as an SCPL family gene based on the common fragments. Sinapoyl-glucose and other cinnamic acid derivatives were found in many studies to be substrates or products of SCPL enzymes^50,60,65–79^. Classification of Solyc12g088250 as an SCPL gene was then confirmed by the conserved motif.

Despite the original discovery in *S. pennellii*, the vast potential in metabolite production, involvement of SCPL genes and their products in biotic stress response^80,81^ and some bioinformatic studies^82,83^, the SCPL genes have not been experimentally characterized in tomato or the closely related *Solanaceae*. Characterized reactions of SCPL genes mostly involve small metabolites. However, in this study we found several large soluble metabolites related to sinapoyl derivatives that, to the best of our knowledge, were previously unknown.

The SCPL gene family is fast expanding^83,84^ and is often found in tandem in genomic clusters. Tissue-specific expression, as seen previously and in our results^57^, contributes to diversification. Furthermore, despite close sequence similarity, genes were found to have diverse substrates^57,72,74^, with only a few amino acid changes sufficient to alter substrate binding abilities^50^. We suggest fragment commonality as a tool to highlight enzyme redundancy and specificity in substrates and products. As we show, features increased by OE of Solyc12g088250 and of Solyc04g077650 share some of the most common fragments, but other fragments are different. Together with tissue-specific expression this can be used to study redundancy and divergence of metabolic gene families. Notably, comparing commonality is possible also when direct feature comparison is not straightforward, for example between experiments run with different UPLC methods.

### Water deficit exposes a purine-permease-associated network of nitrogen-containing metabolites

On chromosome 2, we studied an mQTL with a clear WD-dependent regulation. Both feature and gene expression (eQTL) associations co-located to the same mQTL and were more significant in WD conditions (Fig. 7), indicating that the same polymorphism likely regulates both gene expression and feature levels. The remarkably consistent correlations between feature abundance and Solyc02g071100 gene expression were also seen predominantly in WD, exemplifying the importance of experiments exposing plants to diverse conditions to reveal more of the world of specialized metabolites and their regulation.

Solyc02g071100 is annotated as a purine permease, a member of the purine uptake permeases (PUP) gene family. PUP proteins are membrane-localized proton symporters with multiple transmembrane domains^85^. First identified as adenine transporters, PUPs also transport pyridoxine (vitamin B6)^86^, kinetin, alkaloids including caffeine^87,88^ and nicotine (thereby influencing alkaloid metabolism^89,90^), and benzylisoquinoline alkaloid precursors and opiates^91^. Another important substrate of this gene family is cytokinin^92^, as found in rice^93^ and Arabidopsis^94^. Recently reviewed by Demurtas et al.^95^, the PUP family has been shown to transport different purine ring-containing compounds, with distinct substrate preferences for each gene. All known substrates contain nitrogen, which was also highlighted in our commonality analysis and validated by the ^15^N labeling experiment.

One feature associated with the mQTL was annotated as feruloylputrescine, and two others were similar to it. First found and named in citrus^96^, feruloylputrescine is a hydroxycinnamic acid amide. It shows an organ-specific distribution in several plants^97^ including tomato^98,99^, where it is abundant in developing fruit^100–102^. It is present in leaves and accumulates upon exposure to *Tuta absoluta*^99^ and *Pseudomonas syringae*^98^. A direct link between feruloylputrescine and abiotic stress has rarely been reported in the literature^103,104^, and we found no previous reports of water-stress-related regulation.

Features of the larger subgroup of the mQTL feature group (HCA group B), including feruloylputrescine, were negatively correlated to gene expression and decreased in leaves with Solyc02g071100 OE, and are therefore expected to be the precursors in the metabolism associated with PUP activity. They are characterized by common fragment 222.11, which is not found in the products increased by Solyc02g071100 OE. These products are mostly high-molecular-weight compounds, larger than any compounds described in the literature to be transported by PUP, suggesting that they are formed in the subcellular compartment into which Solyc02g071100 transports the precursors.

Compartmentalization is known to be very important for cellular regulation of metabolite production and accumulation^105,106^. PUP family genes have gene-specific subcellular localization^107^, in either the plasma membrane, tonoplast or endoplasmic reticulum (ER). Like other purine permeases, Solyc02g071100 is predicted to be a transporter with multiple transmembrane domains (Supp. Figure 7D), but its membrane localization is not known. Subcellular localization prediction suggests a membrane localization, either plasma membrane, ER, tonoplast or Golgi (Supp. Figure 7E).

To conclude, although the full mechanism of Solyc02g071100 transport and metabolic influence could not be elucidated by our analysis, commonality analysis and ^15^N labeling point to a role for this gene in the transport of multiple nitrogen-containing compounds larger than previously described.

### Transformation commonality links *Solyc09g091700* to double-bond reduction

On chromosome 9, we found a strong mQTL that was consistent across conditions (Fig. 8). The mQTL was associated with a single SNP with very low LD to others, which is unusual for this population with dense SNPs and LD decay of up to 800 kb^2^. This is probably because the SNP arose relatively recently: it first appears in heirloom varieties, at most hundreds of years ago^2^.

Solyc09g091700 is homologous to alkenal double bond reductase (DBR), or 2-alkenal reductase (AER)^108,109^ family genes. These NADPH-dependent enzymes have been studied in various plant species and share two general motifs AXXGXXG and GXXGXXG^109^, found also in Solyc09g091700. Enzymes of this family catalyze the reduction of the α,β-unsaturated bond of reactive carbonyls^108,110^, nitro-fatty acids (NO2-FAs)^111^ and the reduction of cinnamaldehydes to hydroxydihydrocinnamaldehydes^63,109,112^. The outstanding 2.02 transformation between group B and group A features points to a reduction reaction, matching the proposed function of the gene as a DBR that reduces a C=C double bond by adding H_2_. Consistently, the common fragments of HCA group A features were 2.02 m/z units higher than the common fragments found for B group features (Supp. Figure 9DE).

A tomato AER, encoded by Solyc05g005480, reduces the double bond adjacent to the carbonyl group of Furaneol-like flavor compounds^113^, which have a five-atom ring. In contrast, Solyc09g091700 appears to have a different substrate specificity: its associated features include a cyclohexene or cyclohexane ring. Within this gene family, the enzymes whose substrates are most similar to these metabolites, including a cyclohexene ring, were found in *Nicotiana tabacum*^63^ and peppermint^114^. The *Nt*DBR converts perillaldehyde to dihydroperillaldehyde, similar to the perillyl alcohol and carveol substructures found in the mQTL-associated compounds^63^. Carveol, which is a substructure found in some associated metabolites, is a common volatile^115^. It has an inhibitory effect on food-borne harmful bacteria^116^, a fungus^117^, and harmful insects^118,119^. β-ionone, one of the annotated associated features, is a volatile involved in plant-insect interactions^120^ and gives tomato fruit a pleasant scent^121,122^. Although scarcely mentioned in the literature, the Solyc09g091700 gene expression was up-regulated in a tolerant genotype upon exposure to *Tuta absoluta*^123^, and is co-expressed with insecticidal acyl-sugar acyltransferases^124^. Understanding the role of these compounds and genes in biotic stress responses may help improve plant interactions with insects and pathogens. The proposed function of Solyc09g091700, with its broad range of putative substrates, is a step in this direction.

### Broader uses and outlook

Although commonality analysis was developed here around mQTL feature groups, its underlying principle is broader: features can be grouped by any biologically meaningful criterion and then examined for enriched structural or transformation descriptors. Examples include features that respond significantly to environmental perturbation, or features that change significantly in plants with a specific gene knockout or OE, as shown for the four genes in this work. Features significantly and tightly correlated under any set of variable conditions or present in a specific set of samples, such as an exotic phylogenetic group, can also be subjected to commonality analysis to highlight shared structures. With such grouping, stricter significance thresholds give cleaner groups. With a less stringent group, sub-grouping by fragment similarity by FBMN or another biologically relevant method may be useful. Another benefit of the approach is that it can be applied in retrospect to existing data from existing projects, provided enough MS^2^ spectra are available.

Many mQTLs not examined here contain candidate genes and recurrent descriptors that can be revisited as annotation resources and analytical methods improve. The fragmentation and gene expression information still holds potential for further analysis.

The systematic analysis of mQTL commonality showed that a significantly distinct commonality and outstanding descriptors, either fragments, NLs or transformations, are found in most high-impact mQTLs. The three loci studied here demonstrate how commonality analysis can help glean investigation-worthy descriptors and associations from untargeted metabolite profiling. Testing different descriptor types provides complementary applications: recurrent fragments exposed a shared pathway backbone, recurrent mass differences suggested an enzymatic reaction, and condition-specific fragment patterns guided isotope-based characterization of transporter-associated metabolites. Used together with genetic, transcriptomic, and experimental evidence, commonality analysis provides a flexible route from large, incompletely annotated metabolomic datasets to biologically relevant features, independent of prior annotation.

## Materials and methods

### Plant material for genetic mapping - the TS collection

Seeds of the collection used for GWAS and the genomic variation information for the accessions were kindly provided by Harry Klee (University of Florida). The collection includes 15 *S. pimpinellifolium,* 83 *S. lycopersicum* var. cerasiforme and 300 diverse *S. lycopersicum* tomato varieties including many heirloom and modern cultivars, of which we used 362 accessions. The population was previously genotyped and the population structure has been studied and published^2^. This population was used for mapping of metabolic traits related to fruit flavor^6^.

### Greenhouse experiment setup and treatment regime

Seeds of all accessions of the TS collection were sown in seedling trays and placed in a commercial nursery for germination and establishment. Four weeks after germination, the seedlings were transferred to the experimental greenhouse. The experiment was designed in complete randomized blocks, with one plant of each accession in each block. Three complete randomized blocks of the population were grown in control conditions, akin to commercial growing conditions. Three additional blocks received water deficit (WD) treatment. After two weeks in the greenhouse, irrigation for the WD blocks was gradually reduced until there was a measurable difference in stress measures between WD and control plants (see section below). At this point, WD plants received 12% of the water given to control plants, and this rate was maintained for the remainder of the experiment. Exact watering amounts during the full experiment are reported elsewhere^125^.

### Leaf water content and vegetative production

Leaves from one block of both control and WD were evaluated for water content three weeks after initiation of the WD treatment. The full fourth open leaf was picked and immediately weighed at the greenhouse using a semi-analytical scale. Leaves were then oven dried at 60 °C for 24 h then weighed with an analytical scale. The water content was then calculated by: (fresh weight - dry weight)/fresh weight.

At the end of the growing season, after all fruits were harvested, the stem and leaf biomass was weighed (fresh weight) to estimate vegetative production.

### Leaf sampling for metabolomics and transcriptomics

Leaf sampling was conducted during a four-hour period in the middle of the day (10:00 - 14:00), to reduce variation due to diurnal metabolic cycles. For metabolic analysis, two independent complete blocks from each condition were sampled and analyzed separately to evaluate reproducibility. On each sampling day, a pair of control and WD blocks were sampled to ensure sampling at appropriate hours.

For each sampled plant three distal leaflets from the third, fourth and fifth leaves were pooled together in a vial and immediately flash frozen in liquid nitrogen. Samples were kept completely frozen and stored at -80 °C until extraction.

Samples were ground to a fine powder using a Mixer Mill-MM 200 (Retsch), and 100 mg of frozen powder was weighed for extraction.

### Data processing

All data processing was done in R (version 4.1.3) ^126^ in the RStudio environment (2022.07.1 build 554) using packages as described or custom code available as listed under Data availability.

### Transcriptomic profiling

#### RNA extractions

Of the same frozen leaf powder used for metabolic extraction, 50 mg frozen powder were weighed for RNA extraction. The extraction followed the TRIzol protocol (Merck, TRI Reagent®, Cat no. 93289) with some modifications^127^. Briefly, 1 ml chilled TRIzol was added to each frozen sample, it was then vortexed and placed on ice. After 5 min incubation at room temperature samples were centrifuged 8 minutes at 24000 g at 4 °C. Then chloroform separation was performed twice: the clear upper phase was transferred to a new tube, 0.25 ml chloroform was added, samples were vortexed, incubated 15 minutes at room temperature, centrifuged 8 minutes at 24000 g at 4 °C, and 0.4 ml of the aqueous phase was transferred to a new tube. To that 0.3 ml chilled isopropanol was added and samples were gently mixed and placed at -20 °C overnight. Samples were centrifuged 25 minutes at 24000 g at 4 °C to pellet RNA. The pellet was washed twice with 75% ethanol. Then it was re-suspended in 100 µl double distilled water and 10 µl sodium acetate (3M, pH=5.2) before 0.3 ml chilled absolute ethanol was added and samples were placed at -20 °C overnight. Following, RNA was pelleted and washed twice as described above. Finally, the clean RNA pellet was re-suspended in 30 µl double distilled water. RNA quality was evaluated by agarose gel separation stained with SYBR™ Safe (Invitrogen™, cat no. S33102) and Nanodrop (DeNovix) reading of concentration (260nm) and purity (260/280 and 260/230 >1.8). The concentration of all samples was adjusted to 18 ng/µl for TranSeq library preparation.

#### RNA sequencing by TranSeq

The detailed method of library preparation, sequencing and results processing of the TranSeq method was described by Tzfadia et al.^127^. Briefly, RNA was fragmented using NEB fragmentation buffer (NEB cat. no. E6150S) at 94 °C for 4 minutes. After quick cooling to -20 °C, 3’ ends were pulled down by poly-T beads. Selected poly-A containing fragments were barcoded and converted to cDNA by barcode-poly-T primers for reverse transcription. After barcoding, cDNA samples were pooled before RNAse treatment.

Then cDNA was cleaned from reagents with SPRI beads (Beckman, AMPure XP) and PCR adaptors were ligated to the cDNA. Following a second SPRI bead cleaning, cDNA samples were amplified by PCR. Quality of resulting library was evaluated by Qubit (Invitrogen™, Qubit™ dsDNA HS Cat no. Q32851) for concentration reading and TapeStation (Agilent) for evaluating library size distribution. Six libraries, each pooled from 94 RNA samples (including three quality controls in each library) were sequenced by 100 bp single reads on a NovaSeq machine (Illumina). Reads were aligned to tomato genome version SL2.50 ITAG2.4.

#### TranSeq - filtering, batch effect normalization and eQTL

Raw counts of each gene were normalized using vst in the R package DESeq^128^. Genes absent from more than 90% of the samples were removed. Genes which had median count in the quality control samples lower than 3.4 (empirically determined threshold) were also removed. In order to reduce variation caused by batches of library preparation and sequencing, each gene’s reads were normalized to the median of batches from the same treatment. The resulting normalized reads were evaluated by PCA to assess batch effect and check normalization of quality control samples using pcaMethods^129^.

Normalized gene counts were used for GO term enrichment, feature-gene correlations and expression QTL (eQTL). eQTL were obtained by using gene expression as phenotype for GWAS mapping with the same method described below for mQTLs.

#### Gene Ontology (GO) term enrichment

For each gene expressed in the leaf samples, expression was compared between WD and control samples across the full population. The fold change and significance (Student’s two-tailed t-test) of the change in WD compared to control were calculated in R. The genes with p value lower than Bonferroni corrected α=0.05 and fold change higher than 1.5 were considered up regulated. Conversely, genes with p value lower than Bonferroni corrected α=0.05 and fold change lower than 0.5 were considered down regulated.

The lists of genes either up or down regulated were subjected to GO term enrichment analysis according to the instructions in the Gene Ontology website http://geneontology.org/docs/go-enrichment-analysis using Fisher’s exact test with FDR correction. The full list of expressed genes found in our experiment was used as a reference list.

### Metabolic profiling

#### Metabolomic extraction

Extraction protocol was previously described^130^. In brief: 1 ml pre-cooled (−20 °C) extraction buffer (homogenous methanol/methyl-tert-butyl-ether [1:3] mixture) was added to frozen plant sample. After 10 min incubation at 4 °C and sonication for 10 min in an ultra-sonication bath, 500 μl of methanol/water [1:3] mixture was added. Samples were vigorously mixed by vortex, then centrifuged (5 min, 14,000 g), resulting in separation of the lipophilic and polar phases. Phases were aliquoted for analysis in three chromatography-mass spectrometry platforms. Three hundred microliters of the (upper) lipophilic phase were collected for analysis of lipophilic compounds. Of the polar phase, 300 µl were taken for analysis by LC-MS. All aliquoted samples were fully dried under a vacuum and stored at -80 °C until resuspension. Prior to LC-MS analysis dried samples were re-suspended with 100 μl methanol water (1:4) mixture.

#### LC-MS profiling-polar metabolites

LC-MS data were obtained using a Waters Acquity UPLC system (Waters) fitted with a C18 column, coupled to a Q-Exactive Orbitrap mass spectrometer (Thermo Fisher Scientific). Instrumental settings for quantification in full MS were previously described by Giavalisco *et al.*^131^. In detail, the UPLC separation was performed using an HSS T3 C18 reversed-phase column (100 mm × 2.1 mm × 1.8 μm particles; Waters). The mobile phases were 0.1% formic acid in H2O (BufferA, UPLC grade; BioSolve, http://www.biosolve-chemicals.com) and 0.1% formic acid in acetonitrile (BufferB, UPLC grade; BioSolve). A 2 μl re-suspended sample was loaded per injection. The gradient flow rate was 400 µl/min, and consisted of: 1 minute 99% A, 1-11 minute linear gradient from 99% A to 60% A, 11-13 minutes linear gradient from 60% A to 30% A, 13-15 min linear gradient from 30% to 1% A, hold 1%A until 17 min, 17-17.5 minutes linear gradient from 1% to 99% A, and re-equilibrate the column for 2.5 min (20-min total run time).

The UPLC-separated sample was directly injected into a Q-Exactive mass spectrometer. Each sample was analyzed in both positive and negative polarities via separate, consecutive LC-MS runs. The spectra for quantification of all samples were recorded full-scan and covering a mass range from 100 to 1500 m/z. The resolution was set to 70000, with 3 scans per second, restricting the loading time to 100 ms. The capillary voltage was set to 3.5 kV with a sheath gas flow value of 60 and an auxiliary gas flow of 20 (values are in arbitrary units). The capillary temperature was set to 275°C, whereas the drying gas in the heated electrospray source was set to 350 °C. The skimmer voltage was set to 25 V, whereas the tube lens was set to a value of 130 V. The spectra were recorded from 1 min to 20 min of the UPLC gradients.

#### Chromatogram processing, quantification, filtering, feature consolidation and normalization

The Xcalibur Qual browser (Thermo Fisher Scientific) was used to visualize and explore chromatograms.

REFINER MS 11 (GeneData) software was used for chromatogram processing and quantification. Processing of MS data included peak alignment, removal of isotopic peaks, and chemical noise. Peak identifiers (Cluster ID), retention time (RT) and m/z were obtained along with peak area in each of the samples. The peak area was chosen to quantify the relative abundance of corresponding features in the samples. To reduce the redundancy, all isotopic peaks from the same compound were clustered, and each cluster, represented by its monoisotopic m/z, is referred to as a metabolite feature (hereafter feature). Several features can derive from one compound.

Features with median area intensity lower than 100,000, or which were present in the blank negative controls with median intensity higher than 1,000,000, were removed. Features found in less than 30% of the samples in either condition were also removed.

The resulting list of features was consolidated in order to remove redundant information caused by adducts or possible multiple detected fragments (in-source fragmentation) of the same compound, using a custom R script. The algorithm is based on similar RT and high correlation (r>0.9) between features across samples.

The resulting feature abundance values were log2 transformed and normalized to reduce machine-caused variance using each sample’s median intensity and median machine run batch intensity, using R. The process of leaf sampling, extraction, LC-MS analysis, data processing, filtering and normalization was done separately for two repetitions of the population grown in control and WD conditions.

#### Annotation by 30 representative samples

For detailed annotation of all detected features, 30 representative samples were subjected to multiple methods of annotation. The selected samples were genetically varied from both conditions. Extracted aliquots from the samples were sent for analysis in the lab of Prof. Aharoni for high confidence annotation using WeizMass^47^ and lab in-house libraries. The resulting annotations were compared to ddMS^2^ fragmentation spectra obtained from the same samples by methods described in the next section.

#### LC-MS for ddMS^2^ and list-based ddMS^2^ acquisition

For high-level metabolite annotation, it was crucial to obtain feature-specific fragmentation spectra. For high coverage ddMS^2^ fragmentation of all features, the 30 representative samples were analyzed by LC-MS, using the exact same chromatographic method previously used but with ddMS^2^ Top10 method in the MS. In detail: MS acquisition was during 0.4-16.55 min of the full chromatography run, either polarity was acquired separately. For full MS: resolution - 70000, AGC target - 1e6, max IT - 200 ms, scan range - 100-1500 m/z. For ddMS^2^: resolution - 17500, AGC target - 1e5, max IT - 50 ms, loop count - 10, isolation window - 3.0 m/z, CE - 25, 35 (stepped), spectrum data type - centroid. For dd settings: minimum AGC - 2.5e3, apex trigger - 1 to 4 s, charge exclusion - unassigned, dynamic exclusion - 5.0 s.

Despite the high coverage, not all features mapped by GWAS had ddMS^2^ spectra using the Top10 method. Therefore, features associated with detected mQTLs were reanalyzed in the LC-MS, this time with a ddMS^2^ inclusion list method. A list of features, and their RT and m/z identifiers were included for specific fragmentation. In detail: MS acquisition was during 0-17.58 min of the full chromatography run. For full MS, either polarity was acquired separately: resolution - 70000, AGC target - 3e6, max IT - 200 ms, scan range - 100-1500 m/z. For parallel reaction monitoring (PRM): resolution - 17500, AGC target - 5e5, max IT - 100 ms, isolation window - 3.0 m/z, NCE - 25, 35 (stepped).

#### Compound Discoverer

To maximize annotation methods, the 30 samples were analyzed with Compound Discoverer 3.3 (Thermo Fisher Scientific).

Processing parameters: All spectra between 0.45-16.5 min were acquired. Chromatogram RT was aligned according to the quality control pool sample. For detect compounds: mass tolerance 5 ppm, min peak intensity - 100000, S/N threshold - 1.5, no baseline removal, Zig-Zag index threshold - 0.2, jaggedness threshold - 0.4, modality threshold - 0.9, considered ions- [2M+FA-H]-1; [2M+H]+1; [2M+K]+1; [2M+Na]+1; [2M+NH4]+1; [2M-H]-1; [2M-H+HAc]-1; [M+FA-H]-1; [M+H]+1; [M+H+H2O]+1; [M+K]+1; [M+Na]+1; [M+NH4]+1; [M-H]-1; [M-H+H2O]-1; [M-H+HAc]-1; [M-H-H2O]-1, base ions - [M+H]+1; [M-H]-1. For group compounds: mass tolerance 5 ppm, RT tolerance 0.1 min, align peaks - false, preferred ions: [M+FA-H]-1; [M+H]+1; [M+Na]+1; [M+NH4]+1; [M-H]- 1; [M-H+HAc]-1, area integration - most common ion, peak rating contribution was: area -3, CV -10, FWHMN -5, Zig-Zag index - 5, jaggedness-5, modality threshold - 5, peak rating threshold - 4, number of files -1. For fill gaps: mass tolerance 5 ppm, S/N threshold - 1.5. For QC correction the SERRF algorithm was used: min QC coverage - 50%, max QC area RSD - 30%, max corrected QC area RSD 50%, max # files between QC files - 15, # batches -1, interpolate gap filled QC areas - false.

Detected features were compared to several libraries for annotation: full MS m/z were compared to ChemSpider^132^, an in-house-run central metabolites chemical standards library (MSMLS, Mass Spectrometry Metabolite Library of Standards, Sigma-Aldrich), and literature-derived lists ^20,133–136^ as mass lists, ddMS^2^ fragmentation spectra were compared to mzCloud online database^137^ and multiple acquired databases: Bamba laboratory 598 polar metabolites, Fiehn Vaniya natural product library, MassBank of North America (MoNA) (export 2022), and ReSpect^138^. Priority in annotation was given to in-house-run standards, ddMS^2^ based matching, followed by precursor-ion-based mass-list matching (Supp. Table 3).

#### GNPS and feature-based molecular networking

Global similarity of all detected features was used for extending annotation from annotated features to similar ones. Chromatograms from the 30 representative samples were used, analyzed with the regular full MS method and the Top10 ddMS^2^ method, in either positive or negative mode. First, chromatogram files were converted from raw to mzML format, using ProteoWizard file converter (v. 3.0.21112)^139^. Next, chromatogram and spectra, for each polarity separately, were aligned and processed using MS-DIAL (v. 4.60)^140^, with the following specifications: Ionization type -Soft ionization, Separation type - Chromatography, MS method type - Conventional LC/MS, Data type (MS1) - Centroid, Data type (MS/MS) - Centroid, Ion mode - Positive/Negative, Target omics - Metabolomics. For Data collection: Mass accuracy tolerance for MS1- 0.01 Da, for MS2-0.025 Da, RT range - 0.5-16.5, MS1 mass range - 100-2000 Da, MS/MS mass range 0- 2000 Da, maximum charge - 2. For peak detection: minimum peak height 1000 amplitude, mass slice width - 0.1 Da, smoothing method - Linear weighted moving average, smoothing level - 3 scans, minimum peak width - 5 scans. For MS2 Deconvolution: sigma window value - 0.5, keep isotopic ions until 0.5 Da. For identification: RT tolerance - 0.5 min, accurate mass tolerance for MS1 - 0.01 Da, for MS2 - 0.05 Da, and the rest as default. Adducts considered in positive polarity: [M+H]+, [M+NH4]+, [M+Na]+, [M+CH3OH+H]+ and [M+K]+. Adducts considered in negative polarity: [M-H]-, [M-H2O-H]-, [M+Na-2H]-, [M+FA-H]-, [M+Hac-H]-, [2M-H]-, [2M+FA-H]- and [2M+Hac-H]-. For alignment: reference file was set as the quality control pool sample, RT tolerance - 0.05 min, MS1 tolerance - 0.015 Da, and the rest as default. Alignment results were exported for “GNPS export” in the “mgf” format.

In the Global Natural Products Social Molecular Networking (GNPS)^46^ module for FBMN (v. 28.2) a new project was started for each polarity. The MS-DIAL alignment table, the spectra file in mgf format and all chromatogram files in mzML format were uploaded using an ftp client, WinSCP (v. 5.17.10). FBMN was run with default settings, except Quantification Table Source was set to MS-DIAL, Precursor Ion Mass Tolerance - 0.75 Da, Fragment Ion Mass Tolerance - 0.5 Da.

Results were visualized within the GNPS browser and exported in “Cytoscape data” format for further analysis. Cytoscape (v3.9.1)^141^ was used for further network exploration.

#### CFM-ID in silico fragmentation prediction

Predicted fragmentation patterns of formula proposed by Xcalibur (Thermo Fisher Scientific) were generated using CFM-ID 3.0 online software^142^.

#### Calculation of expected mass from molecular formula

While comparing predicted compositions and library annotations, molecular formulas were used to calculate predicted average molecular weight and monoisotopic mass by R package “OrgMassSpecR”^143^.

#### GWAS mapping with GAPIT

The normalized relative abundance of each feature in each condition was used as the phenotype for mQTL mapping by GWAS. Population single nucleotide polymorphism (SNP) variations were kindly provided by Harry Klee (University of Florida). The GWAS SNP locations and all genome information are based on SL2.50^1^ in SGN browser (https://solgenomics.net/jbrowse_solgenomics/?data=data/json/SL2.50), and gene predictions are based on ITAG2.4, unless otherwise stated.

GWAS analysis was performed using the Genome Association and Prediction Integrated Tool (GAPIT) algorithm implemented in R^144,145^. Generally, the default settings were used, specifically the Mixed Linear Model (MLM) method was used and VanRaden kinship^144,146^ and three principal components were used to control for population structure.

#### mQTL co-location grouping and comparisons

After gathering the results from mapping of all features in all conditions, co-locating mQTLs of all features were identified using a custom script. mQTLs were regarded as one co-locating mQTL if significant SNPs (-log(p)>8.3, Bonferroni correction for α=0.05 for 1.8 million SNPs) were at the most 100 kb apart. The set of features associated with one mQTL is referred to as its mQTL feature group. A feature can belong to more than one mQTL feature group (mQTLs on different chromosomes). Comparison of mQTL feature groups and significance level at each condition was performed.

To evaluate the reproducibility of the mQTL mapping, all features which had significant mapping in either condition of the first control-WD block pair were listed. The corresponding features in the second block pair’s dataset were identified by matching RT and m/z identifiers. The relative abundance of the matched features in the second pair was mapped with GAPIT, as described above, and resulting mQTLs were compared for co-mapping in either condition as described above (but with significance threshold of -log(p)>7).

#### Commonality analysis

The mgf files made by MS-DIAL (v. 4.60)^140^ from all experiments were combined in R, using the “MsBackendMgf” package^147^. The ddMS2 spectra were processed with the “CluMSID” package^148^ and custom scripts. Redundant spectra from features with close RT within 0.05 min and m/z within 0.005 Da, and spectra similarity cosine score above 0.6 were removed. The many fragments with negligible counts are a heavy burden on downstream calculations, therefore, fragments with counts lower than 2% of the highest-count fragment were removed. All spectra with less than 2 fragments were removed, since they could not be used in downstream calculations.

The commonality, the set of descriptor frequencies in a group, was calculated using custom scripts. In brief, all ddMS^2^ spectra in the defined group were assembled. Fragments were binned with an m/z tolerance of 0.005 Da, and the number of detected fragments in all group spectra was calculated. The frequency of a fragment is the proportion of the features with ddMS2 spectra in the group whose spectrum contains it. The frequency of the top 50 fragments in the ddMS^2^ group was plotted as a commonality spectrum. The frequency of each fragment in the commonality of all ddMS^2^ spectra was defined as the expected frequency for further statistical analysis. The G- test^149^ was used to statistically compare commonality of tested groups to the expected frequency. We defined the statistical threshold of significance based on 2,000 randomly sampled ddMS^2^ spectra groups. We found that the G-test power was greatly affected by the size of the group (Supp. Note 1), therefore, subsequent tests were compared to the threshold based on similar group size to determine significance. A group whose commonality differed significantly from the expected commonality by this test is described as having a significantly distinct commonality. Separately, a fragment whose frequency in a group exceeded the mean top frequency of size-matched random groups is described as an outstanding descriptor. This is a descriptive threshold, not a significance test. High-impact mQTLs were defined as mQTLs with enough features to statistically test a descriptor type in one ionization mode: at least 20 ddMS^2^ spectra for fragments, 50 ddMS^2^ spectra for NLs, and 50 features for transformations (Supp. Note 1).

The neutral losses (NLs) for each ddMS^2^ spectrum were calculated as the differences between all fragments, including the full m/z of the precursor ion, rounded to 0.01 Da. All commonality and statistical analysis were then conducted in the same way as described above for fragments. NL annotation was based on a list curated from the literature (Supp. Table 7).

Transformation calculations used the m/z of the precursor ions. A transformation is the m/z difference between the precursor ions of two features in a group. It is a mass difference and does not by itself demonstrate a biochemical conversion. For general transformation commonality the differences between all m/z were binned with tolerance of 0.01 Da and the frequency of each difference was calculated out of the number of non-redundant pairs and plotted. Transformations were compared to calculations from the MetNet R package^31^. For directed transformation commonality, only pairwise m/z differences between opposing groups were included. The expected transformation frequencies were calculated based on m/z differences of all detected precursor ions.

The G-test^149^ was used to statistically compare transformation commonality of tested groups to the expected frequency. We defined the statistical threshold of significance based on 2,000 random permutations of randomly sampled precursor ion m/z values. We found that, here too, the G-test power was greatly affected by the size of the group (Supp. Note 1), therefore, subsequent tests were compared to the threshold based on similar group size to determine significance. Transformation annotation was based on a list curated from the literature (Supp. Table 7).

### Feature and gene expression data analysis

#### Principal component analysis (PCA)

PCA was used to gain initial overview of the samples’ metabolomic profile and evaluate normalization performance, using the R package pcaMethods^129^.

Feature-feature correlations

The features of each mQTL feature group were evaluated by correlations. Correlation matrices were made with the R package corrplot^150^. Features were grouped according to hierarchical clustering and major groups were assigned a letter from A-C. Correlation matrices were made for each condition separately.

#### Feature-gene correlations

For each mQTL, a correlation matrix of the relative abundance of all features in the mQTL feature group to the normalized expression of all genes in the mQTL region was produced. Correlations with p-value <0.05 were plotted. Correlation matrices were made for each condition separately. Correlations with p value <10^-7^ and |r|>0.4 were considered significant.

#### Phenotype single allele interactions

To look at the effect of the mQTL alleles on feature abundance, the genotype of the mQTL lead SNP in each accession was extracted, and the effect of the allele on feature abundance in the samples was tested by Student’s t-test.

#### Linkage disequilibrium (LD)

To assess LD of SNPs in the mQTL region, all SNPs in the mQTL region and extending 5 kb before and after the mQTL were used. Calculations and plotting were performed with the R package LDheatmap^151^.

### Isotope labeling

#### 15N isotope labeling

To provide the seeds the material required for their growth to mature plants, 20 S. lycopersicum seeds were germinated on 1/2 Murashige and Skoog (MS) medium (2.15 g/L) and 3% glucose for 30 d. To label the nitrogen within the plant all nitrogen components within the MS medium were replaced with labeled alternatives: 1650 mg/L Ammonium nitrate (^15^NH_4_ ^15^NO_3_, Sigma-Aldrich: 366528) and 1900 mg/L Potassium nitrate (K^15^NO_3_, Merck: 57654-83-8) in the labeled media. The MS medium was then pH balanced to 5.8 ± 0.1 before adding the glucose and autoclaving for 20 min at 121 °C. The plants grew 30 days in 16-8 light conditions and 24 °C before mature leaves were harvested and snap-frozen using liquid nitrogen for UPLC-MS/MS metabolite analysis.

#### 13C isotope labeling

For 13C isotope labeling *S. lycopersicum* calli were grown in tissue culture from lesions in mature leaves in 0.8% agar MS medium supplemented with 0.1 mg/L indole- 3-acetic acid (IAA) for 30 days in 16-8 light conditions and 24 °C, according to Moscatiello *et al.*^152^. The resulting callus cells were then roughly homogenized using a pipette and grown in media consisting of 4.3 g/L MS medium, 1 μg/mL 1- naphthaleneacetic acid (NAA), and 0.1 μg/mL of both 2,4-dichlorophenoxyacetic acid and kinetin.

To label carbon, the medium was supplemented with 3 g/L of either C_6_H_12_O_6_ glucose (control), or ^13^C_6_H_12_O_6_ (Sigma: 389374). These cell cultures were suspended in a shaking incubator at 25 °C for 32 d before harvesting. The cells were harvested according to a UTHealth Houston cell sample preparation protocol. To process the cells for UPLC-MS/MS the samples were centrifuged for 2 min at 1,500 rpm and the supernatant discarded. 1 mL PBS was then added to the pellets, which were then roughly ground using a pipette and centrifuged for 2 min at 2,000 rpm before removing PBS. Afterwards, samples were snap-frozen using liquid nitrogen and stored at -80 °C until UPLC-MS analysis.

#### Isotope labeled metabolomic detection and analysis

The labeled samples were extracted and prepared and run on the UPLC-MS in the same way as described for leaf samples. Chromatograms were visualized in Xcalibur Qual browser (Thermo Fisher Scientific). For each feature first the expected peak was identified by matching m/z and RT in control samples. Next, the labeled samples were tested for presence of a peak with the expected m/z and absence or reduction of the original unlabeled m/z. Elemental compositions of unannotated features were estimated in Xcalibur. Expected m/z values for either ^13^C or ^15^N labeled compounds were calculated using https://bmrb.io/metabolomics/mol_mass.php.

#### Putative gene annotations

Putative gene annotations were initially taken from ITAG 2.4, and investigated further by BLAST (TAIR, NCBI) to find homologous genes or motifs with known functions.

#### Geneious

All sequence analysis, viewing, alignment, neighbor-joining trees and BLAST searches, were done in Geneious Prime® 2022.1.1 Software, unless otherwise stated.

### Validation by overexpression (OE)

#### Cloning of target genes for OE

Genes of interest were amplified from leaf cDNA using HS TAQ MIX RED (PCR BIOSYSTEMS, London, UK) or PrimeSTAR HS DNA Polymerase (TAKARA, Shiga, Japan) followed by digestion with KpnI and XbaI restriction enzymes (New England Ltd., Hitchin, UK) and ligation using T4 DNA Ligase (New England Ltd., Hitchin, UK) into the multiple cloning site of the *pBTEX* plasmid driven by the CaMV 35S promoter. All primers used in this project are listed in Supp. Table 13. Plasmids were introduced into *Agrobacterium tumefaciens* strain *GV3101* by heat shock. Cotyledons isolated from *S. lycopersicum cv. M82* (for Solyc02g071100 OE) were incubated with *A. tumefaciens*, harboring the *pBTEX::35S::gene* vectors following the protocol described by McCormick^153^, to generate transgenic OE lines. Transformation of Ailsa Craig (for Solyc12g088250 and Solyc04g077650 OE) and Micro-Tom (Solyc09g091700 OE) was performed as described by Ouyang *et al*.^154^.

#### RNA extraction, cDNA synthesis and qPCR

RNA was extracted from leaves using the Total RNA Mini Kit (Plant) from Geneaid (New Taipei, Taiwan). The success of the extraction and the integrity of the RNA was validated using NanoDrop (MaestroNano, New Taipei, Taiwan) and gel electrophoresis. cDNA strands were synthesized using a designated kit (qScript cDNA Synthesis Kit; BIOSYSTEMS, London, UK) and PCR (SimpliAmp™ Thermal Cycler; ThermoFisher Scientific, Waltham, MA, USA), according to the manufacturer’s instructions. Quantitative PCR was performed using the PikoReal 96 Real-Time PCR System (Thermo Scientific, MA, USA) and qPCRBIO SyGreen Blue Mix Hi-ROX kit by PCR Biosystems (London, UK). The different sets of primers used for the amplification of the target genes are listed in (Supp. Table 13). Analysis of the relative gene expression was performed according to the comparative cycle threshold (2−ΔΔCT) method^155^, calibrated using transcript values relative to the endogenous tomato actin gene.

#### OE growing conditions, WD

For metabolomic sampling, WT and OE lines were grown in the greenhouse under optimal irrigation. The fourth true leaf was sampled four weeks after transplanting seedlings to the greenhouse. For Solyc02g071100, OE plants were grown in optimal conditions (100% of previous day transpiration) for two weeks, then irrigation was dropped to 50% of daily transpiration for half of the plants for 10 days. Leaf samples were taken prior to applying the WD regime and 4 and 9 days after, from water-limited and fully irrigated (control) plants.

## Author contributions/Acknowledgment

### Author contributions

L.R. designed and managed experiments, prepared samples, analyzed the data and wrote and revised the manuscript. Y.Y.Z., N.Sa. and A.B-S. prepared overexpression lines and conducted their experiments and sample preparation. T.L-C. and S.W. designed and managed experiments, prepared samples and metabolic data acquisition. C.T. prepared samples and data acquisition for isotope labeling experiments. E.S. prepared samples and developed chromatogram processing and metabolite annotation analysis pipelines and reviewed the manuscript. T.N. and S.P. prepared samples for transcriptomics. I.R., N.Sh., and A.A. acquired data and analyzed samples for metabolite annotation. B.O. and S.D. generated transgenic lines. Y.B. initiated the project, acquired funding, supervised all experiments and wrote and reviewed the manuscript. All authors read and approved the manuscript.

## Acknowledgements

We would like to acknowledge the immense support we received from the Ben- Gurion University’s Ilse Katz Institute for Nanoscale Science & Technology Mass Spectrometry and chromatography unit, Elena Doubijansk, Dr. Mark Karpasas, Dr. Albert Batushansky and Dr. Efrat Forti. This work would not be possible without you. We also thank Dr. Albert Batushansky for critical review of the manuscript. Our thanks are extended to Shai Shaked and the greenhouse workers for the high-quality professional cultivation of the tomato GWAS experiment.

We would also like to thank Prof. Dr. Dirk Walther for discussing the statistical framework for the commonality analysis.

## AI disclosure

AI-assisted language editing was used in preparation of the manuscript. No generative AI was used to write new content.

